# Food acquisition patterns of the three-striped palm squirrel are influenced by interactions: variation in foraging response across an urban-rural gradient

**DOI:** 10.64898/2026.10.29.762696

**Authors:** Rakesh Kumar Muni, Adhithi Balaji, Shivani Jadeja, Nandini Rajamani

## Abstract

Urbanisation significantly alters the habitat and resources accessible to urban animals. Synanthropic animals adapt by exploiting available resources and modulating abundance across the urban habitat matrix. Urban arboreal mammals, such as palm squirrels, encountering altered foraging resources in green spaces and across the matrix, develop strategies to maximise their fitness, including modulating food selection, spatial microhabitat use, and social interactions. We examined whether squirrel abundance affects foraging patterns and whether social interactions at the feeding site regulated individuals’ exploratory tendencies. We conducted cafeteria experiments with *F. palmarum* in rural and urban areas around Tirupati, India and tested for differences in responses to novelty, food selection, and dominance interactions among individuals. We observed a preferential bias towards processed food and approach tendencies were stronger in urban individuals. Urban regions marked with a higher density of squirrels, were associated with higher competitive interactions showing transient social despots of dominance, leading to increased exploration by non-dominant individuals. On the contrary, a lower density of squirrels in rural areas, leads to reduced dominance interactions and subsequent lack of exploration. The potential exploration and consumption of novel resources feeds into the urban loop which enhances population in urban areas, whereas the opposite effect is observed in rural areas. This study highlights how urban commensal organisms, such as squirrels, adopt behaviours to maximise resource utilisation which is modulated by social interactions and can potentially determine the population dynamics in changing landscapes.

## Introduction

Urbanisation has reshaped ecosystems, altering animal behaviour and ecology worldwide. Urban environments often provide non-homogeneous resources; these resources are distributed across the urban areas in a dynamic matrix of green spaces and concretised areas (1). Species which exploit these resources and proliferate well are referred to as synanthropic organisms. Globally, aves, arthropods, and mammals make up the majority of synanthropes and urban commensal wildlife (2). Species modify their behaviour to maximise their survival by modulating dietary breadth (3), changing foraging strategies (4), or shifting spatial microhabitat use (5), including increased activities near garbage or food dumps (6). Urban areas offer an abundance of processed foods, which differ in nutritional content and availability from natural foraging options (7,8). This shift in resource availability can drive behavioural adaptations, potentially altering foraging behaviours and risk tolerance (9). Although all animals are driven by available resources in dynamic landscapes, individuals actively make cognitive decisions to access or obtain them, and employ multiple foraging strategies (10), which can be influenced by organisms’ behavioural decisions and personality traits (10,11). In addition, populations can also show variable responses (12). For instance, urban house sparrows (*Passer domesticus*) exploit human-associated food resources more than non-urban populations (13). Foraging decisions can also be modulated by specific resource-acquisition traits and strategies. For example, fruit bat species exhibit variable foraging strategies, allowing individuals to switch between stationary feeding at high-value sites and acquiring portable items to mitigate risk (4). The efficiency of these strategies is typically associated with the latency to approach and consume, which serves as a proxy for an individual’s risk-reward assessment and familiarity with the resource (14). In urban mosaics, such behavioural variation allows some species to navigate the trade-off between the reward of anthropogenic food and the increased exposure to human-related disturbances (15,16).

Although animals can exhibit a gradient between solitary and social lifestyles, individual foraging decisions can be influenced by population structure and density (17). Population density can influence foraging through two opposing mechanisms. On the one hand, the presence of conspecifics can act as a social facilitator; under the many-eyes hypothesis, shared vigilance reduces perceived predation risk, potentially providing individuals with the safety needed to explore novel food items (18). On the other hand, a high density of individuals at clumped resources leads to intense interference competition. This localised crowding can increase feeding latency for some, while forcing others to diversify their intake (19). Consequently, foraging decisions are dictated by a dynamic interplay of food diversity (14) and conspecific interactions (20,21). In regions where multiple species occupy or compete for the same food resource, the presence of heterospecifics can also influence foraging strategies (18,19).

In high-density urban patches, dominance hierarchies or despotic networks may emerge, where dominant individuals monopolise the most preferred, calorie-dense resources (20). This social restriction imposed by dominant individuals may compel subordinates to overcome their neophobia and sample a wider range of secondary, portable, or unfamiliar anthropogenic foods to survive (21). This may lead to dietary innovation, in which subordinates shift toward accessible or portable resources that can be consumed away from the primary or clustered resources monopolised by dominant individuals, thereby utilising spatial partitioning to avoid dominance and agonistic encounters (22,23).

These dynamics suggest a dichotomy between the "Urban Facilitation Loop" and the "Rural Trap". In the urban loop, high resource density drives more social interaction, which, despite the cost of competition, forces behavioural innovation and dietary breadth. In contrast, rural environments often feature dispersed natural resources that support lower population densities. With minimal interference competition, rural individuals are rarely forced to deviate from primary food choices (24). This may lead to conservatism, where individuals remain highly selective and neophobic, restricted to familiar natural items because the pressure to innovate is absent (25). Thus, behavioural traits and social structures interact with habitat type to drive population-level differences in neophobia and boldness (10,26,27).

Small mammals serve as excellent model systems for studying these dynamics due to their widespread distribution, adaptability, and frequent interactions with human-modified environments (9). Among them, squirrels are highly adaptable species that thrive in both natural and urban habitats (28). Their omnivorous diet, opportunistic foraging behaviour, and ability to exploit diverse food sources make them ideal for examining the effect of urbanisation on dietary preferences and behavioural plasticity (29,30). Cafeteria experiments with urban and rural populations of Eurasian red squirrel (*Sciurus vulgaris*) revealed a preference for high-calorie sugar-based processed foods in urban individuals, suggesting dietary adaptation to anthropogenic resources (31). However, the majority of studies are focused on the western region of the globe, with the larger body of literature focusing on temperate wildlife (32–34). Tropical regions contain greater habitat diversity and a growing urban matrix, where species interaction with human-associated or provisioned food is higher (35). Relatively few studies have focused on tropical groups and have directly compared food selection or foraging strategies between urban and rural populations of the same species, particularly in non-Western urban ecosystems.

Studies suggest food preferences depend on accessibility in generalist species (36). Organisms occupy spaces near anthropogenic food sources to maximise resource uptake (4). These choices are more pronounced in tropical squirrels, where direct consumption strategies predominate, unlike food caching in temperate squirrels (37,38). Temperate squirrels use spatial memory to extract caches and select food items for longer storage (39,40). Tropical squirrels feed on diverse diets, prioritising high-energy foods due to consistent availability (41–43), while exploring patchy food sources. These exploration abilities increase in urban habitats due to inconsistent natural resources (44). Human-associated resources are novel to wild organisms; therefore, explorative populations access alternate food items more readily (45,46). Species that rely on urban areas tend to consume human-associated foods because they are readily available (47).

We investigated how urbanisation shapes resource acquisition by addressing four questions: (1) Do food preferences vary across urban and rural populations? We hypothesise that (H1) urban squirrels will prefer processed items due to familiarity, while rural individuals will remain selective for natural foods. (2) How do the acquisition strategies of the preferred food and latencies to interaction vary? We expect (H2) lower latencies and higher tendencies to approach and acquire preferred processed food in urban areas (3). Are conspecific and social interactions around preferred food resources shaped by dominance? We hypothesise (H3) that higher dominant interactions around preferred food items are due to crowding. If there is high dominance (4), how do displaced subordinates explore secondary resources? We expect (H4) that subordinates will utilise spatial partitioning to feed safely and select less-preferred portable secondary items.

## Methods

### Study species

We studied *Funambulus palmarum* (also known as the Indian palm squirrel, Family: Sciuridae), commonly found in the peninsular region of India and in Sri Lanka (48,49). Majorly arboreal, they forage amongst the canopy but also occupy other strata and have been observed to forage on the ground (50). Their dietary breadth includes nectar from certain flower species, fruits, nuts, and insects, as well as anthropogenic food sources, including discarded food and garbage in urban areas. In India, studies of *F. palmarum* populations in the Western Ghats, where interactions with humans are increasing, have also found them foraging on and eating the fruits of cardamom, cocoa, and coffee plantations (51).

### Study area

We selected 80 sites, 41 in urban areas and 39 in rural areas, from 100m x 100m urban and rural grids in and around the urban agglomeration of Tirupati, Andhra Pradesh, India (13° 37’ 44.6340’’ N and 79° 25’ 28.0056’’ E). The grids were created across an urban gradient, taking factors such as human population density and nighttime activity into account (Swati et al. 2026, for details). The size of the grids was determined with respect to the home ranges of palm squirrels, ensuring that the same animal would not be encountered in multiple adjoining grids (52). The study was carried out over three sampling periods from November 2022 to September 2023. 24 grids each were sampled in November 2022 (12 each in urban and rural areas) and February 2023 (12 each in urban and rural areas), while 17 urban grids and 15 rural grids were sampled in August-September 2023. Within each grid, we selected experiment sites for food experiments. Each of these sites was at least 150-200 m apart to ensure the experiment’s novelty remained intact.

### Field method - site selection and cafeteria experiment

We identified potential sites for the experiments, through preliminary scan sampling, confirming squirrel presence by audible vocalisation or visual sightings. We recorded the behaviour of squirrels, their spatial presence and activity, and associated covariates, including human presence and heterospecifics, within a 50 m vicinity. Sites were selected if they met the following three criteria: (1) at least one squirrel was observed within 1 m of the ground; (2) no human or known non-human threats were present within a 10 m radius; and (3) the site allowed unobstructed access for the squirrels and a clear line of sight for video recording from 10 m away.

We designed a cafeteria experiment that offered seven familiar food items with varying macronutrient compositions (Nigam et al., 2021). We selected food items that were common and observed to be consumed by squirrels in preliminary observations. The cafeteria options included unprocessed items (raw peanuts, raw rice, banana) and processed items (boiled peanuts, peanut butter, cooked rice, banana chips). Food was presented in shallow cups made from dried leaves to avoid any additional effect of the base or plate on the squirrel’s behaviour. The cups were filled with food items at a nearby location and then placed as a cafeteria setup at the experiment site. We set up the experiment 1 m away from a tree or wall where one or more squirrels were observed during our pre-setup scan. The filled cups were arranged in a consistent, alternating pattern of peanut butter (pb), raw rice (rr), boiled rice (br), raw banana (rb), banana chips (bc), raw peanuts (rp), boiled peanuts (bp) in a curved line. This slightly curved arrangement ensured equal visibility to all food types, independent of the squirrel’s entry point. After setting up the experiment, observers (working in pairs) moved 10 m away to prevent any effect of their presence on the squirrels’ behaviours. One observer recorded the interaction around the cafeteria setup with a video camera (Canon Powershot IX350), while the other observer documented the squirrels’ activity. Each experiment lasted 30 minutes (determined after pilot experiments in urban and rural areas and held constant across all experiments), and both observers recorded observations manually using a pair of Nikon Prostaff 3 binoculars.

Two methods of behavioural observation were used: A) A focal scan was initiated if a squirrel approached and interacted with the cafeteria setup. In the case of multiple individuals, the observer chose one individual at random and followed it until it left the setup. After the exit of one squirrel, another individual, if present, then became the new focal individual. B) A scan sample was conducted at regular intervals of five minutes, noting surrounding activities - presence and activity of conspecifics, heterospecifics and predators such as Shikra, cats, dogs, and kites. Human movement and activities within 50 m of the experiment were also noted. These two methods ensured we captured information on squirrel presence in the immediate vicinity of the cafeteria setup, which could influence the choice and time spent interacting with it.

### Data extraction and behavioural video annotation

We extracted data from the behavioural sampling and observations. The number of instances when squirrels (a) approached the cafeteria setup, (b) approached food items (details noted) within the experiment, and (c) fed on food items were noted in detail. The experiments were categorised as ‘successful approach’ or ‘no approach’ based on whether a squirrel approached within 1m of the cafeteria setup during the stipulated 30 minutes or not. We also noted the type of experiment (urban or rural) and the time of the experiment, as well as covariates such as existing food sources in the area and conspecifics consuming or interacting with such food sources. Successful feeding attempts were considered only when squirrels were observed consuming the offered food items, either directly from the food cup or by holding them in their forepaws. Instances where the squirrels only interacted with the set-up, like sniffing or examining the food item(s), but did not consume any item, were recorded as approaches.

We prepared an ethogram (see Table S3.1) based on the activities of the squirrels observed during the pilot experiments. Using the ethogram, we manually decoded video recordings of the experiment using BORIS (Behavioural Observation Research Interactive Software) (53). From the video, the following duration-based factors were recorded: the time to first approach the setup, the time to first feed on an item, the time spent in each behaviour, and the total time spent in the experiment. The number of individuals present and their activities were also noted. The approach diversity (AD) and feeding diversity (FD) scores (0–7) were assigned to each experiment based on video observations, with the AD score reflecting the total diversity of food items approached and the FD score reflecting the number of food items fed upon after an approach. The combined scores reflected the interaction with food in the setup, with a higher diversity score indicating a broader dietary breadth and feeding.

### Statistical analysis

All statistical analyses were performed using the R computing environment (v. 4.4.3, R Core Team, 2025) within the RStudio interface (2025.09.1, Build 401; Posit Software, PBC). We investigated whether the initial food choice differed significantly between urban and rural habitats using a Pearson’s Chi-squared test. However, due to the small sample size of feeding events in rural experiments, which resulted in low expected frequencies, Fisher’s Exact Test was utilised as an alternative. To test whether continuous environmental factors drove specific food choices, we applied the nonparametric Kruskal-Wallis rank-sum test to compare the distributions across the different categories of the first food chosen. All data visualisations were generated using the `ggplot2` (54) and `ggpubr` (55) packages.

We used Generalised Linear Models (GLMs) to account for the hierarchical structure of the duration data. For response variables representing diversity scores (counts), a Poisson error distribution with a log link function was utilised, as indicated by the reported Incidence Rate Ratios (IRRs). We fitted GLMs with a Gamma error distribution and a log link function for latency data, which are positive, continuous, and typically right-skewed. Predictor variables included habitat type (urban vs rural), the numeric count of conspecifics in the vicinity (at 10 m and 50 m), the binary presence of predators, and the binary presence of heterospecifics within 10 meters. To account for local urbanisation intensity, the volume of human movement and the binary presence of vehicular movement were included as covariates.

To investigate the drivers of repetitive selection and return to the food items, we fitted Generalised Linear Mixed Models (GLMMs) with a binomial error distribution and a logit link function using the `lme4` package (56). The response variable was binary (1 = the individual selected the same food type as in the immediate previous bout; 0 = the individual switched to a different food type). The full model included the interaction term between Food Type (7 levels) and Urban/Rural site type, along with the scaled counts of conspecifics in the setup (within 1 m) and in the vicinity (within 10 m) as fixed effects. To account for repeated measures and the hierarchical structure of the data, we included ‘Subject’ nested within Trial ID as random effects. Pairwise comparisons for significant interaction terms were conducted using the `emmeans` package (57), with Tukey’s method for multiple comparisons. Model selection was performed using Akaike’s Information Criterion (AIC), and model fit was assessed using marginal and conditional R² values.

To analyse the factors determining the location of food consumption, we used Multinomial Logistic Regression via the `multinom` function in the `nnet` package (58). This approach was chosen because the response variable, Feeding Location, consisted of three nominal categories: *"At cup"*, *"Within setup"* (inside the cafeteria radius but not at the source), and *"Away"* (retreating outside the cafeteria setup). The category *"At cup"* was set as the reference level. Predictors included food type, social role (dominant/subordinate/neutral), and urban/rural site type. Model coefficients were exponentiated to report Odds Ratios (OR).

To assess the temporal costs of displacement and return after chase, we performed a Time-to-Event (Survival) Analysis using the `survival` (59) and `survminer` packages (60). We defined the "event" as the return to foraging. The time interval (latency) was calculated as the duration (in seconds) between the stop time of a previous foraging bout and the start time of the subsequent bout by the same individual. Intervals where the individual did not return within the observation period were right-censored. Kaplan-Meier survival curves were generated to visualise return rates across social roles (dominant vs subordinate). Differences between groups were tested using the nonparametric log-rank test (`survdiff`).

Dominance networks were reconstructed using the `igraph` package (61). Directed edges (Chaser → Chasee) were defined based on temporal proximity. An edge was created if a "Chaser" event and a "Chasee" event occurred within a 5-second temporal window in the same trial. To prevent pseudoreplication, unique identifiers were generated for every individual in every trial (e.g., `TrialID_SubjectName`), ensuring that networks represented specific experiments rather than a pooled population. We calculated Degree Centrality for each node. Out-Degree (number of chases initiated) was used as a proxy for dominance/despotism, while In-Degree (number of times chased) indicated subordination. The network visualisation was generated using the Fruchterman-Reingold layout algorithm to emphasise social clustering.

## Results

### Food Type Preferences and Diversity

Processed food items were preferred over unprocessed ones across experiments. Peanut butter was the most frequently chosen first food, followed by boiled peanuts, especially in urban experiments. Urban experiments showed a 58-68% preference for peanut butter and a 22% preference for boiled peanuts. Rural squirrels showed a preference similar to that of urban squirrels for peanut butter and boiled rice (66% and 33%, respectively). The difference between the urban and rural experiments was not statistically significant (*χ^2^* = 1.739*, p* = 0.187*, df* = 1).

Dietary breadth differed between environments, with a greater diversity score in urban areas (urban: *M* = 2.5 ± 0.8; rural: *M* = 1.8 ± 0.6; *t* = 4.13, p < 0.001).

### Approach and feeding attempts

The initial comparisons between urban and rural sites revealed significant differences in the squirrels’ interaction with the cafeteria setup. A notably higher proportion of experiments in urban areas recorded squirrels approaching the cafeteria setup compared to rural areas. Specifically, we observed this approach in 22 of 41 urban experiments (53.7%) but in only 9 of 39 rural experiments (23.1%) (*χ^2^* = 6.7542*, df* = 1*, p* = 0.009). Following the initial approach, urban squirrels also showed a significantly greater propensity to make feeding attempts. Among the squirrels that approached the setup, individuals in 18 out of 22 urban experiments (81.8%) proceeded to feed, whereas feeding occurred in only 3 out of 9 rural experiments (33.3%) (*χ^2^* = 8.2312*, df* = 1*, p* = 0.009).

### Latency to first approach and first feeding attempt

While urban squirrels showed quicker engagement with the cafeteria setup, analysis of latency to first approach and feeding attempt showed no significant differences between urban (*W* = 100*, p* = 0.821) and rural populations (*W* = 35*, p* = 0.4508). Though not significant, rural squirrels generally took longer approach times for both behaviours (Urban: *M* = 2.3 ± 0.9 mins; Rural: *M* = 3.1 ± 1.2 mins), and longer time for the first feeding attempt (Urban: *M* = 3.8 ± 1.4 mins; Rural: *M* = 4.7 ± 1.6 mins).

### Factors influencing resource acquisition

Nearby squirrels (within 1 m) (*IRR* = 0.64*, p* < 0.001) and predator presence decreased approach time (*IRR* = 0.69*, p* = 0.01), while conspecifics within 10 meters increased it (*IRR* = 1.21*, p* = 0.003). The trial type variable did not significantly predict approach time (*IRR* = 0.89*, p* = 0.513), despite more urban squirrels approaching. The model showed a strong fit with a marginal *R²* of 0.553 and a conditional *R²* of 0.941.

Urban squirrels took longer to feed after approaching compared to rural ones (*IRR* = 1.67*, p* = 0.039). Presence of squirrels within 1m led to a decrease in feeding latency (*IRR* = 0.64*, p* < 0.001), as did human presence within 10m (*IRR* = 0.69*, p* < 0.001). Higher squirrel abundance increased feeding latency (*IRR* = 1.03*, p* = 0.013). Predator presence decreased time to first feeding (*IRR* = 0.50*, p* < 0.001). The marginal *R²* was 0.544, and the conditional *R²* was 0.95.

Squirrel presence within 1 m significantly influenced dietary breadth (*IRR* = 1.17*, p* = 0.008), indicating that higher local density is associated with greater food diversity. Trial type ( *IRR* = 0.94*, p* = 0.826), abundance (*IRR* = 0.99, *p* = 0.506), dominant interactions, and conspecifics within 10 meters (*IRR* = 0.861*, p* = 0.759) were not significant predictors. The marginal *R²* was 0.542, with a conditional *R²* of 0.923.

### Squirrel abundance and repetitive food interactions

We recorded significantly higher numbers of squirrels in urban than in rural areas (*t* = -3.1815*, p* = 0.0026). The mean squirrel density in urban sites was significantly higher than in rural sites (urban, *M =* 12.4 ± 3.2 squirrels/50 m; rural, *M* = 7.8 ± 2.1 squirrels/50 m; *t* = 7.5601, *p* < 0.001). Dominant interactions, such as chasing or displacing, were more frequent in urban areas (*M* = 4.2 ± 1.5 interactions/trial) than in rural areas (*M* = 2.1 ± 0.9 interactions/trial) (*t* = 7.544, *p* = 0.003). We observed a slight positive relationship between the number of dominance interactions and the total number of squirrels present around the experiment (within 1m) (Estimate = 1.24, SE = 0.28, p > 0.05; Figure 6a). In contrast, the frequency of dominance interactions was a significant positive predictor of the diversity score (Estimate = 2.48, SE = 0.88, p < 0.05; Figure 6b)

The GLMM analysis to determine factors influencing foraging persistence indicated that food type is a primary predictor of repetitive behaviour (*p* < 0.001). Peanut Butter (PB) elicited the highest baseline probability of repetition. In contrast, discrete and portable food items such as Raw Rice (*β* = -2.13*, SE* = 0.59*, p* < 0.001) and Raw Bananas (*β* = -1.97*, SE* = 0.57*, p* < 0.001) were associated with significantly lower odds of repetition. The influence of urbanisation on repetition was context-dependent. Notably, the interaction for Urban X Banana Chips showed a coefficient to be significant and positive (*β* = 2.45*, SE* = 1.20*, p* = 0.041),

### Dominance and Social Interactions

Social rank significantly predicted the likelihood of repetitive feeding. A GLMM showed that dominant individuals (chasers) were significantly more likely to repeat their food choice than neutral individuals (*β* = 0.82*, SE* = 0.38*, p* = 0.032). While subordinate individuals (chasees) did not differ significantly from neutral individuals in their overall probability of repetition (*p* = 0.98), their behavioural sequences revealed a qualitative difference. Dominant individuals exhibited higher repetition for preferred food items (e.g., PB *→* PB transitions > 85%), whereas subordinates showed higher transition probabilities to less preferred food items (e.g., PB *→* Rice).

A Multinomial Logistic Regression confirmed that squirrels employ spatial partitioning to mitigate conflict. Social rank strongly dictated feeding location: subordinate individuals were significantly more likely to move away from the cup to feed compared to dominant individuals (Odds Ratio, *OR* = 3.42*, p* < 0.001). This decision was also mediated by the type of food. Portable items like Boiled Peanuts (*OR* = 60.1*, p* < 0.001) and Raw Peanuts (*OR* = 26.5*, p* < 0.001) strongly predicted away or within setup consumption. Conversely, urban squirrels were significantly less likely to retreat (*OR* = 0.19*, p* < 0.001) compared to rural squirrels.

We analysed the latency to return (the time interval between the end of one foraging bout and the start of the next) to test whether subordinates experience increased return times due to displacement. Contrary to expectations, a Kaplan-Meier Survival Analysis (Log-Rank Test) revealed no significant difference in return latencies across social ranks (*χ^2^* = 3.0*, df* = 2*, p* = 0.2).

Median latencies were uniformly short across all groups: dominant (0.64 s), subordinate (0.64 s), and neutral (0.56 s).

### Social network structure

Social network analysis reconstructed from dyadic interactions (Chaser → Chasee) within each trial revealed a despotic rather than linear dominance structure. The networks were characterised by high centralisation, where a single individual often monopolised aggression. Asymmetry in aggression, as measured by degree centrality scores (Supplementary Table 2), showed a highly skewed Out-Degree (aggression initiated) distribution. In several experiments, a single "despot" accounted for 100% of the chasing events. For example, in one urban trial (DSC_0804), Squirrel A had an Out-Degree of 11 (11 chases initiated) and an In-Degree of 0, while Squirrel 7 had an Out-Degree of 6. Conversely, subordinates were identified by high In-Degree scores with zero aggression output. For instance, in trial MVI_1184, Squirrel D was chased 4 times (In-Degree = 4) without initiating any chases (Out-Degree = 0).

## Discussion

We examined how urbanisation affects the foraging ecology of the *Funambulus palmarum*, focusing on its novel resource use and feeding choices. We show that resource acquisition, food preference and exploitation of anthropogenic foods in urban *F. palmarum* are driven by social interaction and despotic interference competition.

### Food selection and choices in urban and rural areas

We investigated how food selection and foraging strategies in squirrels were influenced by urbanisation using a cafeteria-style experiment, observing variation in resource-acquisition traits across processed and unprocessed food. We found a significant difference in food choices: squirrels in both rural and urban areas preferred processed food. This aligns with our hypothesis that urban individuals prefer high-calorie, readily accessible food items (47,62). However, rural squirrels showing similar food choices contradict previous studies (31,63). Food items are chosen based on nutritional content, but foods that are easily accessible and have lower processing time appear preferable for optimal foraging (63). These patterns show processed food as the first item selected, though unprocessed foods are selected later. This indicates that processed foods are likely incorporated into the diets of commensal animals rather than completely replacing natural foods, as seen in grey langurs (64) and vervet monkeys (65). In addition, our repetition analysis reveals that peanut butter preference was highly repetitive. Once urban squirrels overcome resistance to novelty, they repeatedly approach profitable resources, shifting from exploration to exploitation. Rural squirrels showed lower repetition rates and higher switching, similar to rural coyotes, which test multiple food sources (46).

### Resource acquisition strategies and response latency

Our results showed significant differences between urban and rural experiments in the total proportion of experiments where squirrels approached the experiment, but no difference in latency to approach and feed was observed across trial types. Our results indicate reduced neophobia and increased behavioural boldness in urban populations. But the lower population density in rural areas can also be cited as a reason for fewer instances of approach; however, the presence of squirrels in the vicinity and the lack of approach indicate an avoidance pattern. While more urban squirrels engaged with processed food sources, rural squirrels displayed neophobia towards the cafeteria setup. These patterns align with studies showing that urban animals exhibit reduced neophobia and increased boldness as adaptations to urbanisation (27,66,67). This helps them utilise available food resources, which are often non-uniform and unfamiliar in urban regions. Although the initial approach instances differ between urban and rural areas, the lack of latency differences across repeated approaches indicates that rural squirrels’ approach speed after initial wariness is similar to that of urban individuals. Alternatively, some urban individuals took longer to interact, suggesting the influence of external factors rather than personality traits.

When we compared variables affecting latency, we found that the presence of conspecifics reduced approach latency, whereas distant conspecifics had the opposite effect. This reveals complex risk assessment and social dynamics. Close conspecifics within a meter may act as social facilitators through the dilution effect or social learning, reducing approach latency (68–70). However, studies show conspecifics can negatively influence approach latency through competitive interference (71,72). Some studies suggest dominant individuals approach faster to avoid later competition. Conspecifics at greater distance (10m) increase vigilance and latency as squirrels evaluate the social landscape (73,74). A decreasing approach latency to predator presence suggests a grab-and-go strategy in which bold individuals forage quickly (Cooper et al., 2013; Herberholz et al., 2012). This shows that squirrels integrate social cues and risk into their foraging strategies, highlighting the importance of the microhabitat and immediate social environment in decision-making about resource acquisition. These patterns emerge stronger across heterogeneous landscapes and urban matrices, indicating the importance of responses to social cues in microhabitats over broad habitat classifications in urban settings characterised by higher population densities (75,76).

Across urban matrices and within microhabitats, interacting factors such as the presence of humans, heterospecific non-predatory animals and predators can directly influence foraging decisions. Our results showed no significant effect of human presence on latency to approach or feed, indicating habituation in urban and rural areas. However, squirrels avoided the setup when human numbers were very high around the experiments, suggesting a threshold to tolerance of human presence. Familiarity with humans primarily relates to provisioned food, and areas with higher food abundance correlate with higher squirrel numbers (44). Heterospecifics in the vicinity can act as facilitators or inhibitors, depending on the potential relationship between the interacting groups (77,78). Our results align with studies in which the presence of familiar heterospecifics (such as Jungle Babblers, Mynas, and Bulbuls) facilitated approach, whereas higher numbers of heterospecifics prevented approach, resulting in no average effect on latency (79). Although urban squirrels showed quicker approach, delayed feeding could stem from increased conspecific competition in dense urban settings.

Urban individuals tend to feed rapidly after approach, and higher predator presence reduces latency, indicating prioritisation of rapid food acquisition. These findings show competition and predation as opposing selective pressures shaping feeding behaviour (80,81). Human presence (within 10m) strongly influenced feeding latency (Fig. 5B), suggesting different factors affect foraging decisions. Individuals show higher risk assessment during feeding, where predator and human presence prompt quicker food acquisition. Despite habituation to indirect human disturbances in urban organisms (82), a direct human approach causes delayed response and vigilant behaviours (83). Some studies show human presence can decrease fear response (84) and improve problem-solving (85). These responses depend on the number of humans, gaze direction, and human features (86). Contradictory patterns have been observed in the house finch (*Haemorhous mexicanus*), where urban populations are sometimes more neophobic than rural populations due to the high risk of encounters with traps or novel predators (87). *F. palmarum* appears to align more with mammalian exploiters like macaques, where intelligence and manual dexterity allow them to mitigate the risks of novelty (88,89).

The structural properties of food resources dictated foraging strategies, revealing a clear split between scramble and contest competition. Stationary resources like peanut butter forced consumption from the cup, creating a monopolisable point source favoured by dominant individuals. In contrast, discrete, portable items (Boiled Peanuts, Banana Chips), which were not chosen by the dominant individual, facilitated a grab-and-go strategy for the subordinate individuals, as evidenced by our multinomial model, in which portable foods significantly predicted feeding either away from the setup or within the setup but away from the source cup. This behavioural variation parallels the larder-hoarding vs scatter-hoarding plasticity observed in Eastern Chipmunks (85) and the specific handling-time decisions made by urban Eurasian red squirrels (63).

### Effects of social interaction on the latency and diversity of food consumption

The study recorded significantly higher numbers of squirrels in urban environments, associated with increased dominant interactions. This higher density and competition potentially increase explorative tendencies and dietary diversification. A positive correlation was observed between food diversity and dominant interactions. When preferred food is monopolised by dominant individuals, subordinate squirrels may explore (90,91) novel or less-preferred options, thereby broadening their diet (17,92). This dynamic highlights social dynamics as a key driver of foraging plasticity. Dominance and hierarchy are common in group-living organisms (17), but are not prominent in solitary foragers such as squirrels. Normally, solitary species have fewer interactions due to their sporadic distribution (93); however, interactions can increase in urban areas due to resource clustering, seasonal variation in resources (94), or territorial overlap during breeding (95,96) or with competitor species (97). Provisioned food might instigate collective foraging and increase interactions, including territorial ones (90,91). Further studies on the effects of provisioned food on social dynamics and foraging strategies are needed. Dominance can be sex and age-specific (98,99), with dominant individuals acquiring and guarding preferred food (100), or size-associated, with larger individuals dominating smaller ones (17,101). We could only observe unmarked squirrel populations and lacked specific age and gender data; hence, drivers of dominance cannot be specified. Further studies with marked individuals and monitored populations can enhance understanding of social dynamics in foraging.

Our results provided strong evidence for social facilitation. The presence of conspecifics near the setup (within a meter) significantly decreased the latency to approach and, crucially, increased the diversity of food items consumed. This supports the local enhancement hypothesis, which holds that the presence of feeding individuals signals safety and resource availability (102). In high-density urban sites, an initial approach by a bold individual is observed and then followed by other squirrels, creating a cascade of approaches. This mechanism explains the higher dietary breadth observed in urban areas compared to rural sites. In rural areas, lower densities result in fewer opportunities for social information transfer. Without the safety-in-numbers cue, rural squirrels are likely to remain conservative. This phenomenon is well-documented in flocking birds; for instance, large flocks of House Sparrows (*Passer domesticus*) solve foraging puzzles faster than solitary individuals (103). However, unlike simple flocking, our results show that competition drives diversity. When the primary resource (Peanut Butter) is occupied by a dominant, the social pressure forces subordinates to test alternative, previously ignored foods (e.g., Raw Bananas), thereby expanding the population’s realised niche.

Thus, competition and dominance act as significant drivers of foraging flexibility and dietary diversification in urban settings, both of which are density-dependent behavioural adaptations. This highlights the importance of social dynamics, rather than just resource availability, in shaping urban adaptation (104–106). Higher densities in parks, institutions, and urban green spaces are an important consideration in development and sustainable management strategies. Food provisioning regulations and feeding zones are usually part of such strategies, but consideration of population densities and the potential for increased stress or conflict due to high intraspecific competition in such areas is also essential for developing sustainable strategies. This finding reinforces the idea that immediate social competition within the microhabitat is a critical factor in determining foraging choices, resource acquisitions, and dietary diversity in *F. palmarum*, potentially more influential than the broad urban or rural classification.

### Social hierarchy and the effect of dominance on foraging strategy

The social network analysis revealed a despotic structure in urban populations, characterised by high centralisation, in which specific chasers monopolise resources. However, contrary to the expectation that dominance would strictly limit subordinate foraging, our survival analysis showed no significant difference in return latency between dominant and subordinate individuals. We observed that subordinates have developed highly efficient coping mechanisms to coexist with despots. Rather than fleeing the area (which would result in long latencies), subordinates employ spatial partitioning. Our location analysis confirmed that subordinates are significantly more likely to feed away or within setup rather than at the cup. They effectively trade the highest-quality resource (Peanut Butter) for lower-quality or portable resources (Rice, Peanuts) that allow them to maintain a high feeding tempo without engaging in direct conflict. This is consistent with the producer-scrounger dynamics seen in chacma Baboons (*Macaca radiata*), where subordinates exploit the leftovers of the dominant’s forage (107). The identified networks were generally sparse and disconnected, confirming that aggression is not a group-wide phenomenon but is typically driven by specific dominant individuals defending a monopolisable resource (Peanut Butter) against specific targets. The urban environment supports high densities not by eliminating aggression, but by providing enough resource heterogeneity (clumped vs scattered) to allow subordinates to forage peripherally without exclusion (108).

### The urban facilitation loop and rural trap hypotheses

Integrating our findings, we propose that *F. palmarum* adaptation is driven by a density-dependent behavioural feedback loop, which manifests differently in urban and rural contexts.

The Urban Loop: Anthropogenic food subsidies support higher population densities. High density increases encounter rates and social interaction. Frequent interactions facilitate social learning and reduce neophobia via local enhancement. Squirrels explore and exploit a wider range of novel foods and expand their dietary niche. Increased energy intake supports even higher densities. This positive feedback loop creates a population of bold, explorative, diverse foragers who are constantly innovating. Similar loops have been proposed for the success of invasive Argentine ants (*Linepithema humile*) (109,110) and urban pigeons (*Columba livia*) (111), where density breeds efficiency.

The rural trap hypothesis predicts that low resource availability in rural areas limits density (112). Low rates of social interaction reduce opportunities for individuals to exchange social information. Neophobia remains high among individuals as a survival mechanism against predation and threats. This leads to restricted diet breadth, including familiar, natural items. Shifts in such resources, or the introduction of new resources due to anthropogenic changes, are not explored or utilised effectively by these individuals, thereby preventing the population from growing. This hypothesis offers a mechanistic explanation for why some species adapt while others fail. Species that are strictly solitary or highly territorial (e.g., Mountain Lions) cannot leverage the urban facilitation loop because high density leads to lethal conflict rather than social learning (113). *F. palmarum*, being socially tolerant and capable of spatial partitioning, can enter the loop, turning the challenge of crowding into an opportunity for information transfer, exploration and foraging flexibility.

## Conclusion

Our study indicates that the squirrels deploy different strategies to acquire resources in urban and rural regions. Our findings reveal a duality: while urban squirrels prefer highly palatable, processed foods, their overall diet diversity in the experimental setup also increases compared to that of rural squirrels. This suggests that the abundance of diverse anthropogenic food sources allows for both a strong preference for certain high-value items and the exploration of a broader dietary repertoire, driven by social interaction, facilitation and competition. The results underscore adaptive, context-dependent decision-making in urban wildlife, reflecting foraging strategies in response to dynamic environmental risks and opportunities.

This is a crucial finding because it suggests that while urbanisation creates the conditions for higher densities and thus more social interactions, it is the immediate social environment (at a microhabitat scale) that directly drives dietary diversification, rather than the macro-level urban classification itself. The urban environment provides the stage (high density), but the social play on that stage (conspecific interactions) dictates the specific foraging outcome (dietary breadth). This implies that social learning or local competitive pressures within a very small radius can be powerful modifiers of individual foraging behaviour.

**Figure 1.**
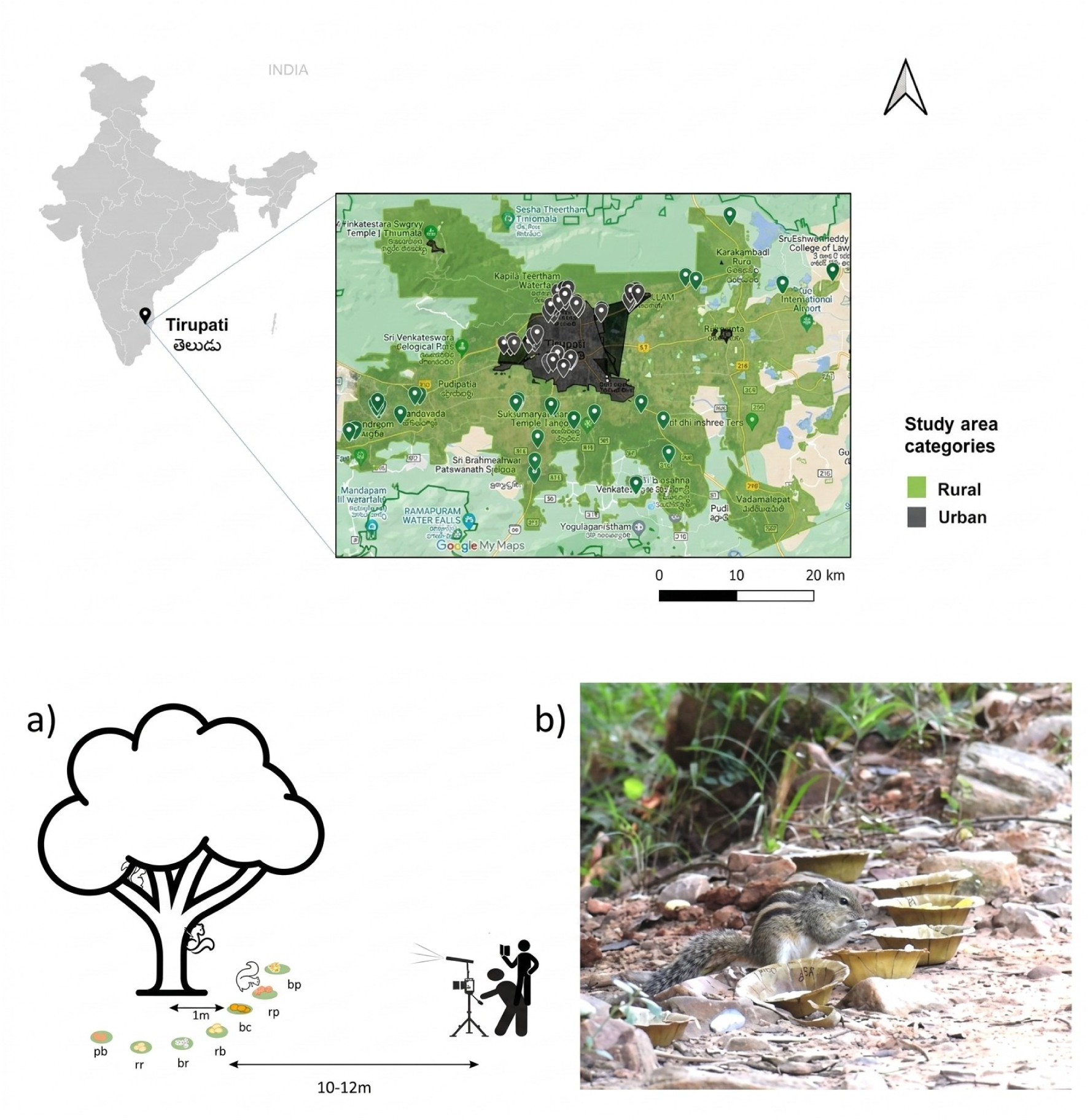
Map of study sites and the experiment sites are marked across urban and rural areas of Tirupati (top). (a) Sketch of food arrangement and cafeteria setup with 7 food items. (b) The three-striped palm squirrel is consuming and interacting with the cafeteria experiment.

**Figure 2.**
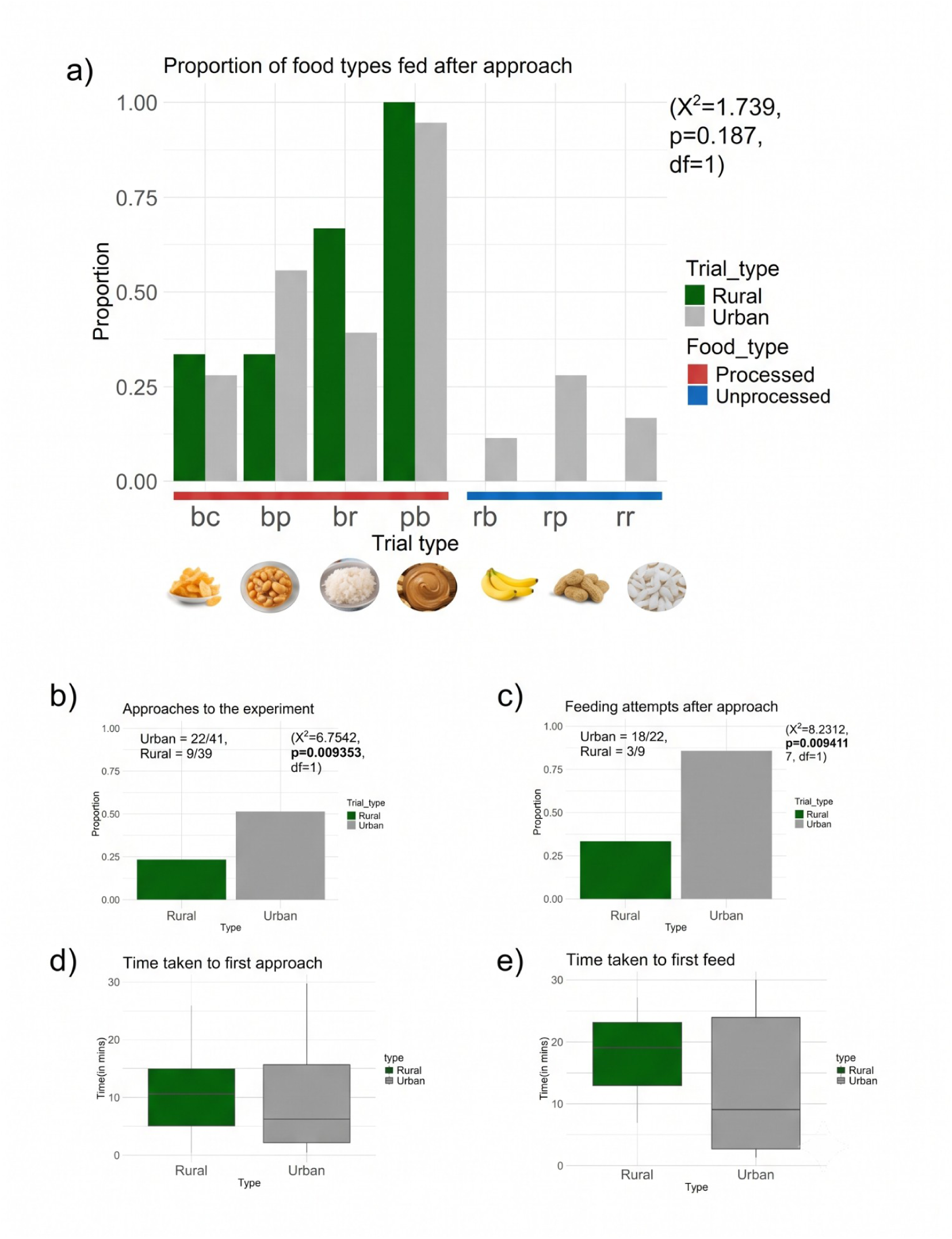
Proportion of food types approached across rural (green) and urban (grey) areas; Processed food included bc – Banana chips, bp – Boiled peanuts, br – Boiled rice, pb – Peanut butter, while non-processed food were rb – Raw banana, rp – raw peanut, rr – raw rice. Panel B: Bar plots for proportion of (a) approaches to experiment, (b) feeding attempts after approach, and Box plots for (c) latency to first approach, (d) latency to first feeding attempt, across rural and urban experiments.

**Figure 3.**
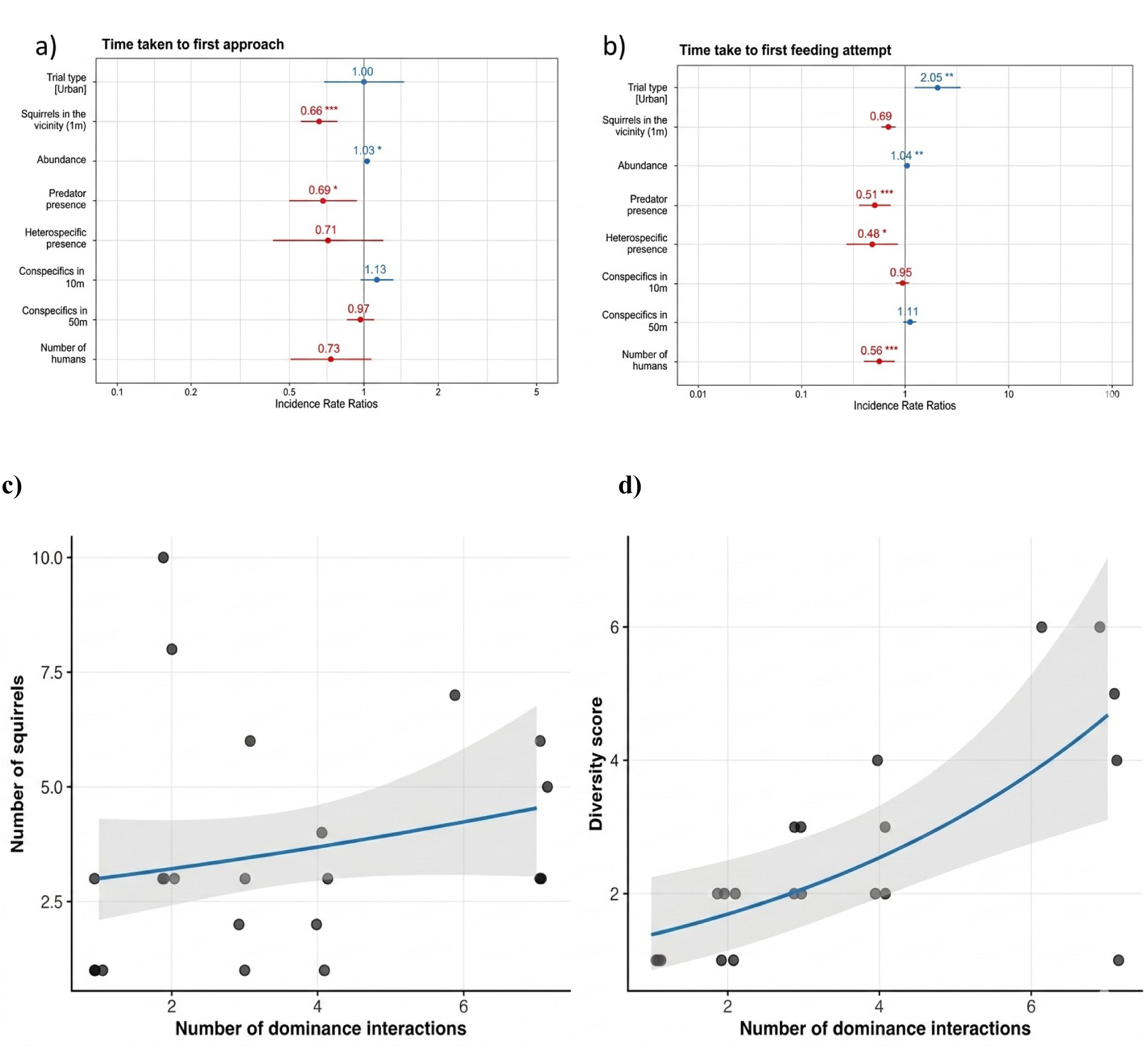
GLM Models and incidence rate ratios for factors influencing **(a)** latency to first approach and **(b)** first feeding attempt. Figure 6. Relationships between dominance interactions, squirrel abundance, and diversity scores during active encounters. **(c)** The total number of squirrels present as a function of the number of dominance interactions. **(d)** The diversity score as a function of the number of dominance interactions. In both panels, solid blue lines denote the predicted fits from Poisson Generalised Linear Models (GLMs), and grey shaded regions represent the 95% confidence intervals. Data points represent individual observational experiments; experiments with zero dominance interactions were excluded from this analysis.

**Figure 4.**
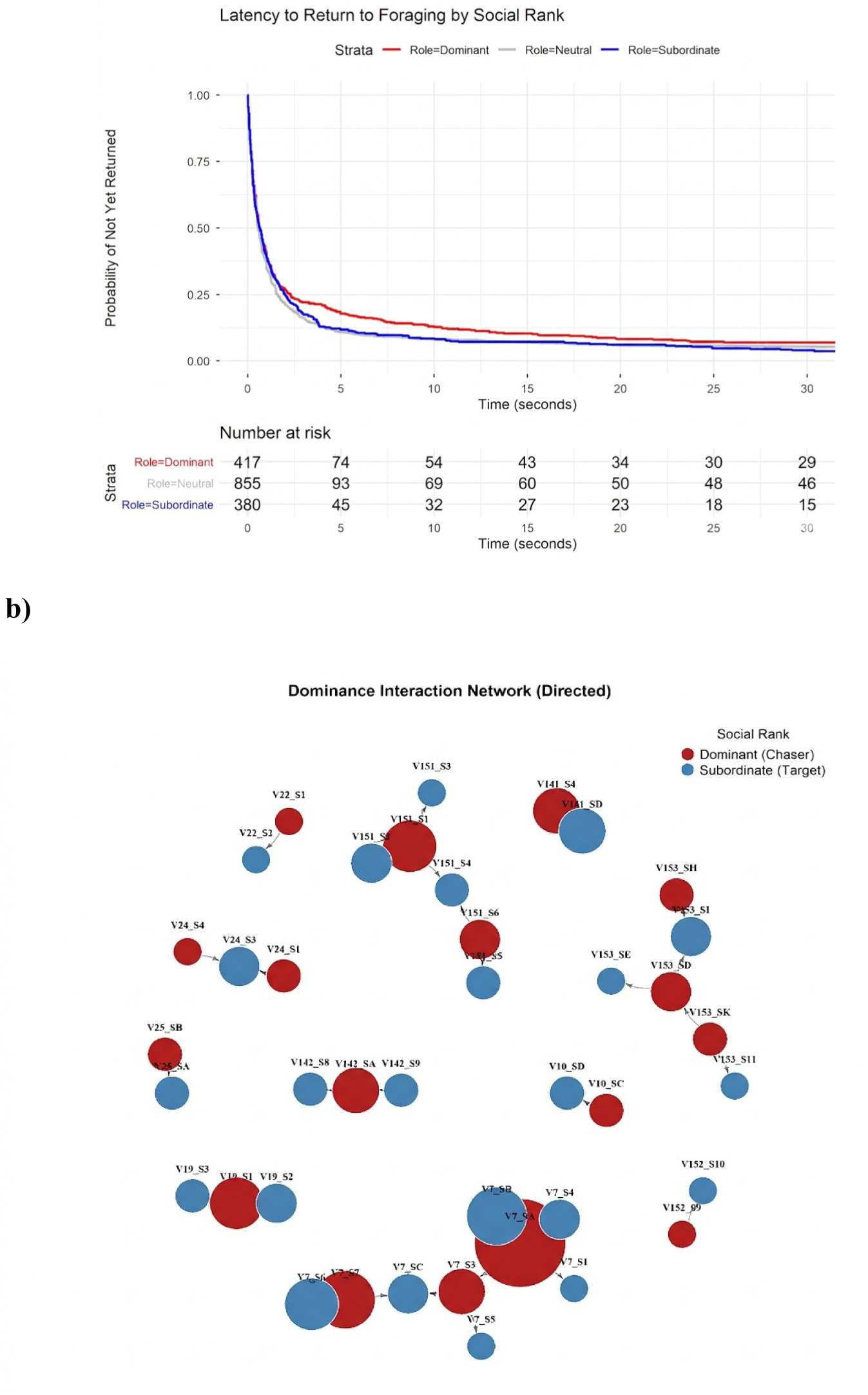
Survivorship curve for latency to return of squirrels for foraging by social rank (top). Social network graph interpreting dominance direction and interaction. The size of the circles is proportional to the number of interactions, and arrows represent the direction of dominance interaction. Each subcluster represents interacting individuals and is named [Video subcluster_Squirrel ID].

## Acknowledgements

This work was supported by intramural funds from IISER Tirupati to NR, support funds from Krea University to AB and by RM’s CSIR-UGC Fellowship (Contingency funds - Award letter no. 191620203010). We thank the Andhra Pradesh Forest Department (Permit number: <u>Rc.no</u>.8549/2019/WL-3) for research permits. We thank Qazi Hammad Moueen and Siva Sankari G for assistance with fieldwork.

## Declaration of generative AI use

The authors report generative AI was not used in preparation of this manuscript.

## References

1. Turner BL II. Land system architecture for urban sustainability: new directions for land system science illustrated by application to the urban heat island problem. J Land Use Sci [Internet]. 2016 Nov 1 [cited 2026 May 29];11(6):689–97. Available from: 10.1080/1747423X.2016.1241315

2. Faeth SH, Bang C, Saari S. Urban biodiversity: patterns and mechanisms: Urban biodiversity. Ann N Y Acad Sci [Internet]. 2011 Mar 1 [cited 2025 July 27];1223(1):69–81. Available from: 10.1111/j.1749-6632.2010.05925.x

3. Anders JL, Mychajliw AM, Moustafa MAM, Mohamed WMA, Hayakawa T, Nakao R, et al. Dietary niche breadth influences the effects of urbanization on the gut microbiota of sympatric rodents. Ecol Evol [Internet]. 2022 Sept 1 [cited 2025 July 27];12(9):e9216. Available from: 10.1002/ece3.9216

4. Egert-Berg K, Handel M, Goldshtein A, Eitan O, Borissov I, Yovel Y. Fruit bats adjust their foraging strategies to urban environments to diversify their diet. BMC Biol [Internet]. 2021 June 16;19(1):123. Available from: 10.1186/s12915-021-01060-x

5. Ancillotto L, Viviano A, Mori E, Bani L, Orioli V, Tirozzi P, et al. Microhabitat features shape small mammal urban assemblages across Italian cities. Sci Total Environ [Internet]. 2025 Aug 1 [cited 2025 July 27];988(179826):179826. Available from: 10.1016/j.scitotenv.2025.179826

6. Plaza PI, Lambertucci SA. How are garbage dumps impacting vertebrate demography, health, and conservation? Glob Ecol Conserv [Internet]. 2017 Oct 1 [cited 2025 July 27];12:9–20. Available from: 10.1016/j.gecco.2017.08.002

7. Carpenter M, Savage AM. Nutrient availability in urban food waste: carbohydrate bias in the Philadelphia–Camden urban matrix. J Urban Ecol [Internet]. 2021 Jan 22;7(1):juab012. Available from: 10.1093/jue/juab012

8. Coogan SCP, Raubenheimer D, Zantis SP, Machovsky-Capuska GE. Multidimensional nutritional ecology and urban birds. Ecosphere [Internet]. 2018 Apr;9(4):e02177. Available from: 10.1002/ecs2.2177

9. Ritzel K, Gallo T. Behavior Change in Urban Mammals: A Systematic Review. Frontiers in Ecology and Evolution [Internet]. 2020;8. Available from: https://www.frontiersin.org/article/10.3389/fevo.2020.576665

10. McElreath R, Luttbeg B, Fogarty SP, Brodin T, Sih A. Evolution of animal personalities. Nature [Internet]. 2007 Nov 8;450(7167):E5; discussion E5-6. Available from: 10.1038/nature06326

11. Sih A, Ferrari MCO, Harris DJ. Evolution and behavioural responses to human-induced rapid environmental change. Evol Appl [Internet]. 2011 Mar;4(2):367–87. Available from: 10.1111/j.1752-4571.2010.00166.x

12. Humphrey JE, Haslem A, Bennett AF. Avoid, adapt or exploit: Re-visiting bird responses to urbanization using a novel landscape approach. Glob Ecol Conserv [Internet]. 2023 Dec 1 [cited 2025 July 27];48(e02735):e02735. Available from: 10.1016/j.gecco.2023.e02735

13. Bókony V, Kulcsár A, Tóth Z, Liker A. Personality traits and behavioral syndromes in differently urbanized populations of house sparrows (Passer domesticus). PLoS One [Internet]. 2012 May 4 [cited 2025 June 16];7(5):e36639. Available from: 10.1371/journal.pone.0036639

14. Prat Y, Yovel Y. Decision making in foraging bats. Curr Opin Neurobiol [Internet]. 2020 Feb 1 [cited 2025 July 27];60:169–75. Available from: 10.1016/j.conb.2019.12.006

15. Aravind PS, Ganguli SP, D’Souza S, Rajamani N. Exotic timber plantations as nesting sites for the endemic Indian Giant Squirrel Ratufa indica in the Palani Hills, Western Ghats, India. Glob Ecol Conserv [Internet]. 2025 Dec 1 [cited 2026 Oct 28];64(e03916):e03916. Available from: 10.1016/j.gecco.2025.e03916

16. Sol D, Lapiedra O, González-Lagos C. Behavioural adjustments for a life in the city. Anim Behav [Internet]. 2013 May 1;85(5):1101–12. Available from: https://www.sciencedirect.com/science/article/pii/S0003347213000511

17. Rands SA, Pettifor RA, Rowcliffe JM, Cowlishaw G. Social foraging and dominance relationships: the effects of socially mediated interference. Behav Ecol Sociobiol [Internet]. 2006 Aug 19 [cited 2025 July 28];60(4):572–81. Available from: 10.1007/s00265-006-0202-4

18. Caraco T, Martindale S, Pulliam HR. Avian flocking in the presence of a predator. Nature [Internet]. 1980 June [cited 2026 Mar 17];285(5764):400–1. Available from: https://www.nature.com/articles/285400a0

19. Bibi N, Wang H. The effect of individual personality on novel food preference in a social context in the cinereous tit (Parus cinereus). Behaviour [Internet]. 2023 Jan 25 [cited 2026 Sept 30];160(3–4):1–14. Available from: 10.1163/1568539X-bja10205

20. Gomez-Melara JL, Acosta-Naranjo R, Castellano-Navarro A, Beltrán Francés V, Caicoya AL, MacIntosh AJJ, et al. Dominance style predicts differences in food retrieval strategies. Sci Rep [Internet]. 2021 Feb 1 [cited 2026 Mar 16];11(1):2726. Available from: 10.1038/s41598-021-82198-0

21. Senior AM, Charleston MA, Lihoreau M, Buhl C, Raubenheimer D, Simpson SJ. Evolving nutritional strategies in the presence of competition: a geometric agent-based model. PLoS Comput Biol [Internet]. 2015 Mar;11(3):e1004111. Available from: https://pmc.ncbi.nlm.nih.gov/articles/PMC4376532/

22. Kikuchi DW. Who innovates? Abundance of novel and familiar food changes which animals are most persistent. Proc Biol Sci [Internet]. 2024 Jan 31 [cited 2026 Mar 17];291(2015):20231936. Available from: 10.1098/rspb.2023.1936

23. Smith MM, Erb JD, Pauli JN. Reciprocated competition between two forest carnivores drives dietary specialization. J Anim Ecol [Internet]. 2023 Sept;92(9):1695–706. Available from: 10.1111/1365-2656.13962

24. Romero-Vidal P, Luna Á, Fernández-Gómez L, Navarro J, Palma A, Tella JL, et al. Intraspecific competition and individual behaviour but not urbanization affect the dietary patterns of a generalist avian predator. Sci Rep [Internet]. 2023 June 24 [cited 2026 Mar 17];13(1):10255. Available from: 10.1038/s41598-023-37026-y

25. Smith DF, Litvaitis JA. Foraging strategies of sympatric lagomorphs: Implications for differential success in fragmented landscapes. Canadian Journal of Zoology [Internet]. 2011 Feb 15 [cited 2026 Mar 17];78(12):2134–41. Available from: 10.1139/z00-160

26. Burkhard TT, Dochtermann NA, Charmantier A. Global meta-analysis reveals urban-associated behavioural differences among wild populations. J Anim Ecol [Internet]. 2026 Sept 1 [cited 2026 Sept 30];95(9):1539–57. Available from: 10.1111/1365-2656.70269

27. Lee VE, Thornton A. Animal cognition in an urbanised world. Front Ecol Evol [Internet]. 2021 Mar 4 [cited 2025 July 27];9:633947. Available from: 10.3389/fevo.2021.633947

28. Swati U, Aravind PS, Muni R, Devi AM, D’Souza S, Rithika, et al. Urban green cover drives high palm squirrel abundance in South India [Internet]. Social Science Research Network. 2025 [cited 2026 Oct 25]. Available from: https://papers.ssrn.com/abstract=5886053

29. Tranquillo C, Bisi F, Wauters LA, Preatoni D, Martinoli A, Santicchia F. The impact of urbanisation on chipmunks, arboreal and flying squirrels: a global systematic review. Mamm Rev. 2023 Dec;

30. Bertolino S. Animal trade and non-indigenous species introduction: the world-wide spread of squirrels. Divers Distrib. 2009 July;15(4):701–8.

31. Wist B, Stolter C, Dausmann KH. Sugar addicted in the city: impact of urbanisation on food choice and diet composition of the Eurasian red squirrel (Sciurus vulgaris). J Urban Ecol [Internet]. 2022 Jan 13 [cited 2023 Aug 4];8(1). Available from: https://academic.oup.com/jue/article-pdf/8/1/juac012/48416315/juac012.pdf

32. Koprowski JL, Nandini R. Global hotspots and knowledge gaps for tree and flying squirrels. Curr Sci. 2008;

33. McDonald RI, Mansur AV, Ascensão F, Colbert M, Crossman K, Elmqvist T, et al. Research gaps in knowledge of the impact of urban growth on biodiversity. Nat Sustain [Internet]. 2019 Dec 9 [cited 2025 July 27];3(1):16–24. Available from: 10.1038/s41893-019-0436-6

34. Trimble MJ, van Aarde RJ. Geographical and taxonomic biases in research on biodiversity in human-modified landscapes. Ecosphere [Internet]. 2012 Dec 1 [cited 2025 July 27];3(12):art119. Available from: 10.1890/ES12-00299.1

35. Erinjery JJ, Kumar S, Kumara HN, Mohan K, Dhananjaya T, Sundararaj P, et al. Losing its ground: A case study of fast declining populations of a “least-concern” species, the bonnet macaque (Macaca radiata). PLoS One [Internet]. 2017 Aug 23;12(8):e0182140. Available from: 10.1371/journal.pone.0182140

36. Reher S, Dausmann KH, Warnecke L, Turner JM. Food availability affects habitat use of Eurasian red squirrels (Sciurus vulgaris) in a semi-urban environment. J Mammal [Internet]. 2016 Dec 5 [cited 2021 June 17];97(6):1543–54. Available from: https://academic.oup.com/jmammal/article-pdf/97/6/1543/8286156/gyw105.pdf

37. Hao H, Jiang C, Wu J, Zong C, Zeng D, Alimubike N. The effects of scatter-hoarding rodents on Mongolian oak (Quercus mongolica) acorn dispersal and seedling distribution in urban woodland. Urban Ecosyst [Internet]. 2017 Dec;20(6):1277–83. Available from: 10.1007/s11252-017-0679-9

38. Thompson MJ, Morand-Ferron J. Food caching in city birds: urbanization and exploration do not predict spatial memory in scatter hoarders. Anim Cogn [Internet]. 2019 June 3;22(5):743–56. Available from: 10.1007/s10071-019-01271-4

39. Delgado MM, Jacobs LF. Caching for where and what: evidence for a mnemonic strategy in a scatter-hoarder. R Soc Open Sci [Internet]. 2017 Sept;4(9):170958. Available from: 10.1098/rsos.170958

40. Hopewell L, Leaver L, Lea S. Effects of competition and food availability on travel time in scatter-hoarding gray squirrels (Sciurus carolinensis). Behavioral Ecology [Internet]. 2008 Nov 1;19:1143–9. Available from: 10.1093/BEHECO/ARN095

41. Aravind PS, Joe G, Dhanesh P, Nandini R. Food habits of the Dusky-striped Squirrel Funambulus sublineatus (Mammalia: Rodentia: Sciuridae). J Threat Taxa [Internet]. 2021 Feb 27 [cited 2026 Oct 28];13(2):17827–31. Available from: 10.11609/jott.6202.13.2.17827-17831

42. Emmons LH. Ecology and resource partitioning among nine species of African Rain Forest squirrels. Ecol Monogr [Internet]. 1980 Mar;50(1):31–54. Available from: 10.2307/2937245

43. Palmer RR, Koprowski JL. Feeding behavior and activity patterns of Amazon red squirrels. Mammalia [Internet]. 2014 Jan 1;78(3). Available from: 10.1515/mammalia-2013-0083

44. Rimbach R, Butler G, Gupte PR, Jäger J, Parker C, Pontzer H. Gray squirrels consume anthropogenic food waste most often during winter. Mamm Biol [Internet]. 2023 [cited 2025 July 27];103(1):69–81. Available from: 10.1007/s42991-022-00326-3

45. Federspiel IG, Garland A, Guez D, Bugnyar T, Healy SD, Güntürkün O, et al. Adjusting foraging strategies: a comparison of rural and urban common mynas (Acridotheres tristis). Anim Cogn [Internet]. 2017 Jan [cited 2025 July 27];20(1):65–74. Available from: 10.1007/s10071-016-1045-7

46. Breck S, Poessel SA, Mahoney P, Young J. The intrepid urban coyote: a comparison of bold and exploratory behavior in coyotes from urban and rural environments. Sci Rep [Internet]. 2019 Feb 14;9. Available from: 10.1038/s41598-019-38543-5

47. Wong BBM, Candolin U. Behavioral responses to changing environments. Behav Ecol [Internet]. 2015 May 1 [cited 2024 Jan 30];26(3):665–73. Available from: https://academic.oup.com/beheco/article-pdf/26/3/665/7720923/aru183.pdf

48. Swati U, D’Souza S, Aravind PS, Muni RK, Rajamani N. A comprehensive database of squirrel distribution and occurrence in South Asia. Biodivers Data J. 2023 Oct;11:e109946.

49. Datta A, Nandini R. Sciurids. In: Manjrekar AJ & editor. Mammals of South Asia. 2015. p. Chapter: 64.

50. Balasubramanian P. Some notes on the fruits, seeds and nectar consumed by three striped palm squirrel Funambulus palmarum at Point Calimere Wildlife Sanctuary, Tamil Nadu. J Bombay Nat Hist Soc [Internet]. 1995;92:256–8. Available from: https://www.biodiversitylibrary.org/part/156175

51. Chakravarthy AK, Thyagaraj NE, Kumar L, Girish A. Crop raiding and management of Funambulus palmarum in cardamom (Elettaria cardamomum) plantations of Western Ghats of Karnataka, south India. Current Science [Internet]. 2008 Dec 7 [cited 2025 July 27];95(7):907–11. Available from: https://www.jstor.org/stable/24103189

52. Prakash I. Palm Squirrel (Funambulus Palmarum, Linnaeus) in Australia. J Bombay Nat Hist Soc [Internet]. 1967;64:108–108. Available from: https://www.biodiversitylibrary.org/part/152896

53. Friard O, Gamba M. BORIS: a free, versatile open-source event-logging software for video/audio coding and live observations. Methods Ecol Evol [Internet]. 2016 Nov 1 [cited 2025 July 27];7(11):1325–30. Available from: 10.1111/2041-210X.12584

54. Wickham H. Ggplot2: Elegant graphics for data analysis [Internet]. 2nd ed. Cham, Switzerland: Springer International Publishing; 2016. 260 p. (Use R!). Available from: 10.1007/978-3-319-24277-4

55. Kassambara A. Comprehensive R Archive Network (CRAN). 2026 [cited 2026 June 9]. “ggplot2” Based Publication Ready Plots [R package ggpubr version 0.6.3]. Available from: https://CRAN.R-project.org/package=ggpubr

56. Bates D, Mächler M, Bolker B, Walker S. Fitting linear mixed-effects models Usinglme4. J Stat Softw [Internet]. 2015 Oct 7 [cited 2025 July 27];67(1):1–48. Available from: 10.18637/jss.v067.i01

57. Searle SR, Speed FM, Milliken GA. Population marginal means in the linear model: An alternative to least squares means. Am Stat [Internet]. 1980 Nov [cited 2026 May 29];34(4):216–21. Available from: 10.1080/00031305.1980.10483031

58. Venables WN, Ripley BD. Modern applied statistics with S [Internet]. New York, NY: Springer; 2010 [cited 2026 May 24]. 498 p. (Statistics and Computing). Available from: 10.1007/978-0-387-21706-2

59. Therneau TM. Comprehensive R Archive Network (CRAN). 2026 [cited 2026 Sept 30]. Survival Analysis [R package survival version 3.8-12]. Available from: https://CRAN.R-project.org/package=survival

60. Kassambara A, Kosinski M, Biecek P, Fabian S. Comprehensive R Archive Network (CRAN). 2026 [cited 2026 Sept 30]. Drawing Survival Curves using “ggplot2” [R package survminer version 0.5.2]. Available from: https://CRAN.R-project.org/package=survminer

61. Csárdi G, Nepusz T, Traag V, Horvát S, Zanini F, Noom D, et al. Comprehensive R Archive Network (CRAN). 2026 [cited 2026 Sept 30]. Network Analysis and Visualization [R package igraph version 2.3.3]. Available from: https://CRAN.R-project.org/package=igraph

62. Méndez A, Montalvo T, Aymí R, Carmona M, Figuerola J, Navarro J. Adapting to urban ecosystems: unravelling the foraging ecology of an opportunistic predator living in cities. Urban Ecosyst [Internet]. 2020 Oct;23(5):1117–26. Available from: 10.1007/s11252-020-00995-3

63. Kostrzewa A, Krauze-Gryz D. The choice and handling of supplemental food by red squirrels in an urban park. Behav Processes [Internet]. 2020 Sept;178:104153. Available from: 10.1016/j.beproc.2020.104153

64. Dasgupta D, Banerjee A, Karar R, Banerjee D, Mitra S, Sardar P, et al. Altered food habits? Understanding the feeding preference of free-ranging gray langurs within an urban settlement. Front Psychol [Internet]. 2021 Apr 26 [cited 2025 July 27];12:649027. Available from: 10.3389/fpsyg.2021.649027

65. Thatcher HR, Downs CT, Koyama NF. Understanding foraging flexibility in urban vervet monkeys, Chlorocebus pygerythrus, for the benefit of human-wildlife coexistence. Urban Ecosyst [Internet]. 2020 Dec 24 [cited 2025 July 27];23(6):1349–57. Available from: 10.1007/s11252-020-01014-1

66. Greggor AL, Jolles JW, Thornton A, Clayton NS. Seasonal changes in neophobia and its consistency in rooks: the effect of novelty type and dominance position. Anim Behav [Internet]. 2016 Nov;121:11–20. Available from: 10.1016/j.anbehav.2016.08.010

67. Quesada J, Chavez-Zichinelli CA, Garcia-Arroyo M, Yeh PJ, Guevara R, Izquierdo-Palma J, et al. Bold or shy? Examining the risk-taking behavior and neophobia of invasive and non-invasive house sparrows. Animal biodiversity and conservation [Internet]. 2022;45(1):97–106. Available from: https://researchportal.helsinki.fi/files/175714466/ABC_45_1_pp_97_106.pdf

68. Charalabidis A, Dechaume-Moncharmont FX, Petit S, Bohan DA. Risk of predation makes foragers less choosy about their food. PLoS One [Internet]. 2017 Nov 9;12(11):e0187167. Available from: 10.1371/journal.pone.0187167

69. St. Lawrence S, Rojas-Ferrer I, Morand-Ferron J. Does the presence of a conspecific increase or decrease fear? Neophobia and habituation in zebra finches. Ethology [Internet]. 2021 Dec;127(12):1033–41. Available from: 10.1111/eth.13224

70. Voelkl B, Schrauf C, Huber L. Social contact influences the response of infant marmosets towards novel food. Anim Behav [Internet]. 2006 Aug 1 [cited 2025 July 27];72(2):365–72. Available from: 10.1016/j.anbehav.2005.10.013

71. Andrade RS, Cerveira AM, Mathias M da L, Varela SAM. Interaction time with conspecifics induces food preference or aversion in the wild Algerian mouse. Behav Processes [Internet]. 2023 Sept;211(104927):104927. Available from: 10.1016/j.beproc.2023.104927

72. Nettle D, Cronin KA, Bateson M. Responses of chimpanzees to cues of conspecific observation. Anim Behav [Internet]. 2013 Sept;86(3):595–602. Available from: 10.1016/j.anbehav.2013.06.015

73. Drakeley M, Lapiedra O, Kolbe JJ. Predation risk perception, food density and conspecific cues shape foraging decisions in a tropical lizard. PLoS One [Internet]. 2015 Sept 18 [cited 2025 July 27];10(9):e0138016. Available from: 10.1371/journal.pone.0138016

74. Stöwe M, Kotrschal K. Behavioural phenotypes may determine whether social context facilitates or delays novel object exploration in ravens (Corvus corax). J Ornithol [Internet]. 2007 Dec;148(S2):179–84. Available from: 10.1007/s10336-007-0145-1

75. Skórka P, Lenda M, Martyka R, Tworek S. The use of metapopulation and optimal foraging theories to predict movement and foraging decisions of mobile animals in heterogeneous landscapes. Landsc Ecol [Internet]. 2009 May;24(5):599–609. Available from: 10.1007/s10980-009-9333-0

76. Fardell LL, Nano CEM, Pavey CR, Dickman CR. Small prey animal foraging behaviors in landscapes of fear: Effects of predator presence and human activity along an urban disturbance gradient. Front Ecol Evol [Internet]. 2022 Mar 30 [cited 2025 July 27];10:805891. Available from: 10.3389/fevo.2022.805891

77. Kanishka AM, MacGregor C, Evans MJ, Dexter N, Dickman CR, Robinson NM, et al. The effect of competitor presence on the foraging decisions of small mammals. Anim Behav [Internet]. 2025 June 1 [cited 2025 July 27];224(123195):123195. Available from: 10.1016/j.anbehav.2025.123195

78. Chen CC, Liao CC, Walther BA. Influence of foraging guild on heterospecific preference of association in mixed-species bird flocks: distinguishing potential benefits for attendant species. J Ornithol [Internet]. 2024 July 29 [cited 2026 Oct 1];166(1):145–56. Available from: 10.1007/s10336-024-02198-5

79. Miller R, Garcia-Pelegrin E, Danby E. Neophobia and innovation in Critically Endangered Bali myna, Leucopsar rothschildi. R Soc Open Sci [Internet]. 2022 July 20 [cited 2025 July 27];9(7):211781. Available from: 10.1098/rsos.211781

80. Dantzer B, McAdam AG, Humphries MM, Lane JE, Boutin S. Decoupling the effects of food and density on life-history plasticity of wild animals using field experiments: Insights from the steward who sits in the shadow of its tail, the North American red squirrel. J Anim Ecol [Internet]. 2020 Nov;89(11):2397–414. Available from: 10.1111/1365-2656.13341

81. Lowry H, Lill A, Wong BBM. Behavioural responses of wildlife to urban environments. Biol Rev Camb Philos Soc [Internet]. 2013 Aug;88(3):537–49. Available from: 10.1111/brv.12012

82. Vincze E, Papp S, Preiszner B, Seress G, Bókony V, Liker A. Habituation to human disturbance is faster in urban than rural house sparrows. Behav Ecol [Internet]. 2016 Apr 1 [cited 2025 July 27];27(5):1304–13. Available from: 10.1093/beheco/arw047

83. Blumstein DT, Fernández-juricic E, Zollner PA, Garity SC. Inter-specific variation in avian responses to human disturbance: Predicting wildlife responses to tourism. J Appl Ecol [Internet]. 2005 Oct 1 [cited 2025 July 27];42(5):943–53. Available from: 10.1111/j.1365-2664.2005.01071.x

84. Ortiz-Jimenez CA, Conroy SZ, Person ES, DeCuir J, Gall GEC, Sih A, et al. Human presence shifts the landscape of fear for a free-living mammal. Ecology [Internet]. 2025 Jan 1 [cited 2025 July 27];106(1):e4499. Available from: 10.1002/ecy.4499

85. Chow PKY, Uchida K, Koizumi I. “Ripple effects” of urban environmental characteristics on cognitive performances in Eurasian red squirrels. J Anim Ecol [Internet]. 2024 Aug 1 [cited 2025 July 27];93(8):1078–96. Available from: 10.1111/1365-2656.14126

86. Goumas M, Lee VE, Boogert NJ, Kelley LA, Thornton A. The Role of Animal Cognition in Human-Wildlife Interactions. Front Psychol [Internet]. 2020 Nov 4;11:589978. Available from: 10.3389/fpsyg.2020.589978

87. Hutton P, Lendvai ÁZ, Németh J, McGraw KJ. Urban house finches are more resistant to the effects of artificial light at night. Sci Total Environ [Internet]. 2024 Oct 10 [cited 2026 Mar 18];946(174525):174525. Available from: 10.1016/j.scitotenv.2024.174525

88. Jaman MF, Huffman MA. The effect of urban and rural habitats and resource type on activity budgets of commensal rhesus macaques (Macaca mulatta) in Bangladesh. Primates [Internet]. 2013 Jan;54(1):49–59. Available from: https://idp.springer.com/authorize/casa?redirect_uri=https://link.springer.com/article/10.1007/s10329-012-0330-6&casa_token=BinBacTJP4kAAAAA:usFuNhxby_yrGMH6g5eMy9nLpImviDWm4xqA7zXOaSHsWR_ymMKTifv8LxWAHAJJac1tAohd-qollaTThA

89. Marty PR, Balasubramaniam KN, Kaburu SSK, Hubbard J, Beisner B, Bliss-Moreau E, et al. Individuals in urban dwelling primate species face unequal benefits associated with living in an anthropogenic environment. Primates [Internet]. 2020 Mar;61(2):249–55. Available from: https://idp.springer.com/authorize/casa?redirect_uri=https://link.springer.com/article/10.1007/s10329-019-00775-4&casa_token=u7f2SPVpOIIAAAAA:SMqgcVLkUpn1i_zpD775r3XPPAKUth4CFptrleXN7PIiEwGdh03F_H_NBkKnySom9kckU1zI0T-0lDWbwg

90. Støstad HN, Aldwinckle P, Allan A, Arnold KE. Foraging on human-derived foods by urban bird species. Bird Study [Internet]. 2017 Apr 3 [cited 2025 July 28];64(2):178–86. Available from: 10.1080/00063657.2017.1311836

91. Wilson CA, Hernandez S, Weil JN, Ezenwa VO, Altizer S, Hall RJ. Human-provided food increases aggregation but does not change activity budgets in an urban wading bird. J Urban Ecol [Internet]. 2024 Feb 27 [cited 2025 July 28];10(1):juae017. Available from: 10.1093/jue/juae017

92. Pollack L, Culshaw-Maurer M, Sih A. Social dominance influences individual susceptibility to an evolutionary trap in mosquitofish. Ecol Appl [Internet]. 2025 Jan 1 [cited 2025 July 28];35(1):e3081. Available from: 10.1002/eap.3081

93. Makuya L, Schradin C. Costs and benefits of solitary living in mammals. J Zool (1987) [Internet]. 2024 May 1 [cited 2025 July 28];323(1):9–18. Available from: 10.1111/jzo.13145

94. Li X, Wang X, Lu J, Li L, Li D, Xing X, et al. Eating more and fighting less: Social foraging is a potential advantage for successful expansion of bird source populations. Biology (Basel) [Internet]. 2022 Oct 12 [cited 2025 July 28];11(10):1496. Available from: 10.3390/biology11101496

95. Clutton-Brock T. Social evolution in mammals. Science [Internet]. 2021 Sept 17 [cited 2025 July 28];373(6561):eabc9699. Available from: 10.1126/science.abc9699

96. Lane JE, Boutin S, Gunn MR, Coltman DW. Sexually selected behaviour: red squirrel males search for reproductive success. J Anim Ecol [Internet]. 2009 Mar 1 [cited 2025 July 28];78(2):296–304. Available from: 10.1111/j.1365-2656.2008.01502.x

97. Muni RK, Rajamani N. Resource acquisition strategies among palm squirrel species across sympatry and allopatry. In Prep. 2026;

98. Enoksson B. Age- and sex-related differences in dominance and foraging behaviour of nuthatches Sitta europaea. Anim Behav [Internet]. 1988 Feb 1 [cited 2025 July 28];36(1):231–8. Available from: 10.1016/S0003-3472(88)80266-5

99. Lewis S, Phillips RA, Burthe SJ, Wanless S, Daunt F. Contrasting responses of male and female foraging effort to year-round wind conditions. J Anim Ecol [Internet]. 2015 Nov 1 [cited 2025 July 28];84(6):1490–6. Available from: 10.1111/1365-2656.12419

100. Dorning J, Harris S. Dominance, gender, and season influence food patch use in a group-living, solitary foraging canid. Behav Ecol [Internet]. 2017 Sept 1;28(5):1302–13. Available from: 10.1093/beheco/arx092

101. Alatalo RV, Moreno J. Body size, interspecific interactions, and use of foraging sites in tits (paridae). Ecology [Internet]. 1987 Dec;68(6):1773–7. Available from: 10.2307/1939868

102. Galef BG. Imitation and local enhancement: detrimental effects of consensus definitions on analyses of social learning in animals. Behav Processes [Internet]. 2013 Nov 1 [cited 2026 Mar 18];100:123–30. Available from: 10.1016/j.beproc.2013.07.026

103. Liker A, Barta Z. The effects of dominance on social foraging tactic use in house sparrows. Behaviour [Internet]. 2002 Jan 1 [cited 2026 May 25];139(8):1061–76. Available from: 10.1163/15685390260337903

104. McKinney ML. Urbanization, biodiversity, and conservation. Bioscience [Internet]. 2002 Oct 1 [cited 2025 July 27];52(10):883. Available from: https://academic.oup.com/bioscience/article/52/10/883/354714?login=true

105. Ditchkoff SS, Saalfeld ST, Gibson CJ. Animal behavior in urban ecosystems: Modifications due to human-induced stress. Urban Ecosyst [Internet]. 2006 Jan [cited 2025 July 22];9(1):5–12. Available from: 10.1007/s11252-006-3262-3

106. Miranda AC. Mechanisms of behavioural change in urban animals: The role of microevolution and phenotypic plasticity. In: Ecology and Conservation of Birds in Urban Environments [Internet]. Cham: Springer International Publishing; 2017 [cited 2025 July 27]. p. 113–32. Available from: 10.1007/978-3-319-43314-1_7

107. King AJ, Isaac NJB, Cowlishaw G. Ecological, social, and reproductive factors shape producer–scrounger dynamics in baboons. Behav Ecol [Internet]. 2009 July 13 [cited 2026 Mar 18];20(5):1039–49. Available from: 10.1093/beheco/arp095

108. Macdonald DW, Johnson DDP. Patchwork planet: the resource dispersion hypothesis, society, and the ecology of life: Patchwork planet: resource dispersion and the ecology of life. J Zool (1987) [Internet]. 2015 Feb 1 [cited 2026 May 25];295(2):75–107. Available from: 10.1111/jzo.12202

109. Calcaterra LA, Chifflet L, Fernández MB, Pirk GI, Werenkraut V, Sánchez-Restrepo AF. From Pampas to Patagonia: Human-modified environments drive the spread of the Argentine ant beyond its climatic limits. Diversity (Basel) [Internet]. 2025 Sept 24;17(10):667. Available from: https://www.mdpi.com/1424-2818/17/10/667

110. Menke SB, Booth W, Dunn RR, Schal C, Vargo EL, Silverman J. Is it easy to be urban? Convergent success in urban habitats among lineages of a widespread native ant. PLoS One [Internet]. 2010 Feb 12;5(2):e9194. Available from: https://journals.plos.org/plosone/article?id=10.1371/journal.pone.0009194

111. Szulkin M, Munshi-South J, Charmantier A. Urban evolutionary biology [Internet]. Szulkin M, Munshi-South J, Charmantier A, editors. London, England: Oxford University Press; 2020 [cited 2026 Mar 18]. 320 p. Available from: https://books.google.com/books/about/Urban_Evolutionary_Biology.html?id=rJfgDwAAQBAJ

112. Battin J. When good animals love bad habitats: Ecological traps and the conservation of animal populations: Ecological traps. Conserv Biol [Internet]. 2004 Dec 1 [cited 2026 June 1];18(6):1482–91. Available from: 10.1111/j.1523-1739.2004.00417.x

113. Benson JF, Sikich JA, Riley SPD. Individual and population level resource selection patterns of mountain lions preying on mule deer along an urban-wildland gradient. PLoS One [Internet]. 2016 July 13;11(7):e0158006. Available from: https://pmc.ncbi.nlm.nih.gov/articles/PMC4943639/

