## Supplementary for "Food acquisition patterns of the three-striped palm squirrel are influenced by interactions: variation in foraging response across an urban-rural gradient"

**Supplementary- Food choice paper**

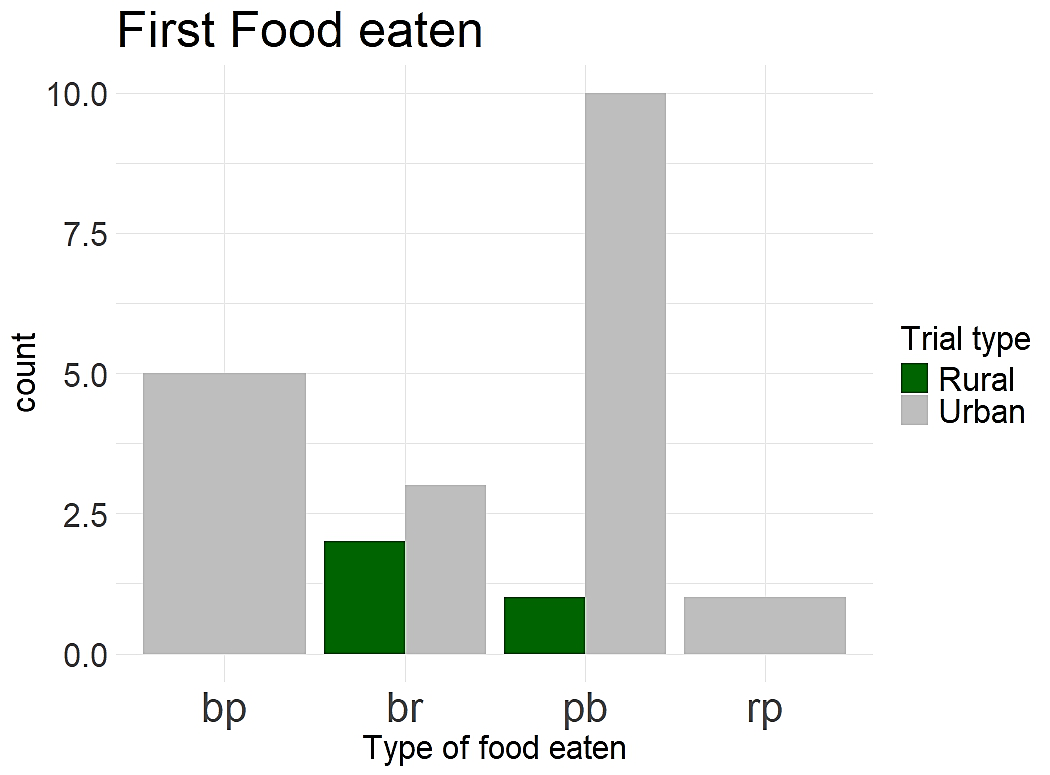

Fig. S1. Frequency of the food item eaten in the first attempt across rural and urban trials

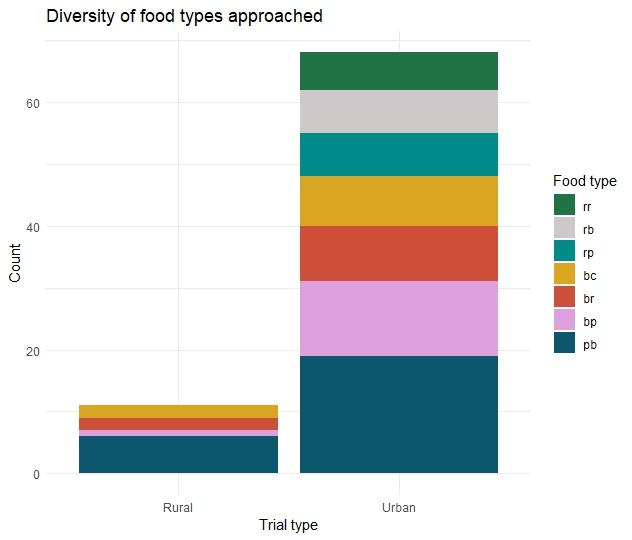

Fig. S2. Diversity of the food items approached across rural and urban trials

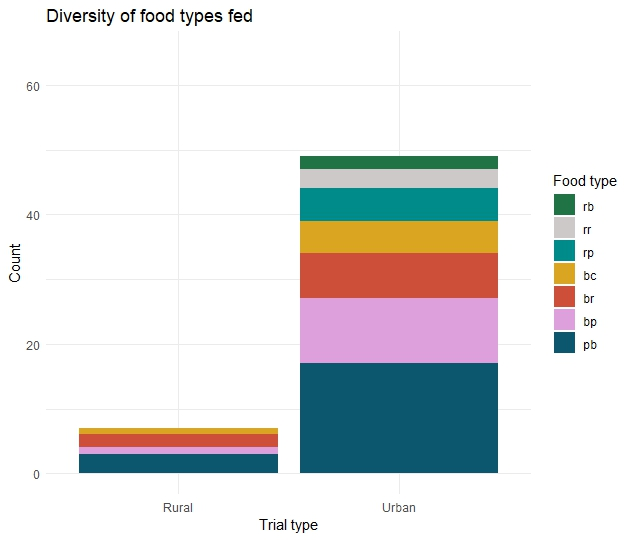

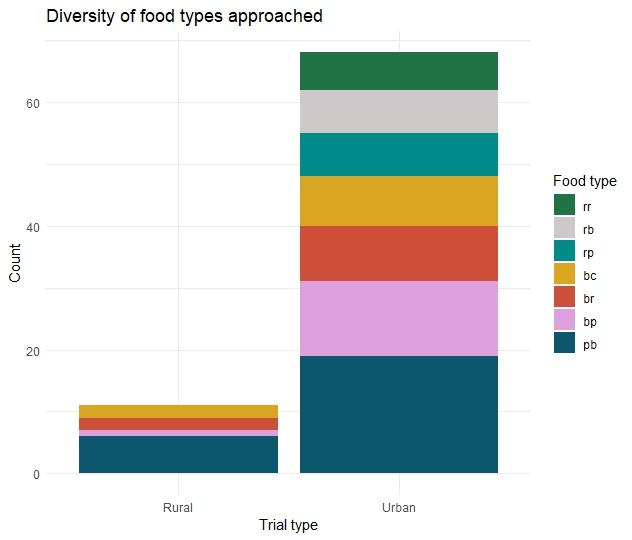

Fig. S3. Diversity of the food items fed upon by squirrels across rural and urban trials

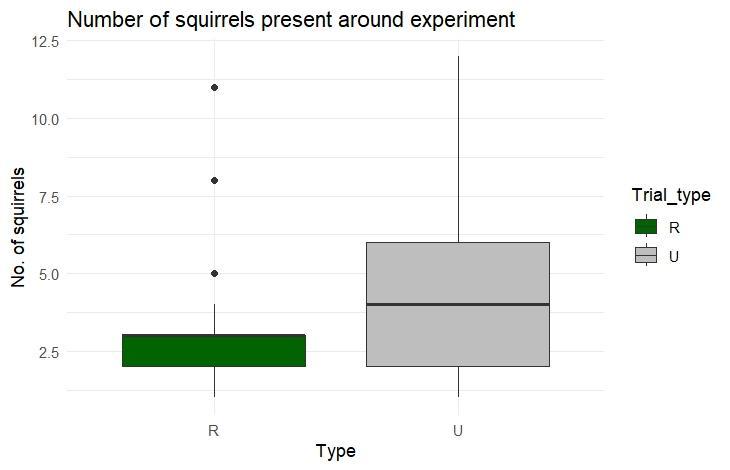

t = -3.1815,

**p-value = 0.0026**

Fig. S4. Density of squirrels in the vicinity(1m) of the experiment across rural and urban trials

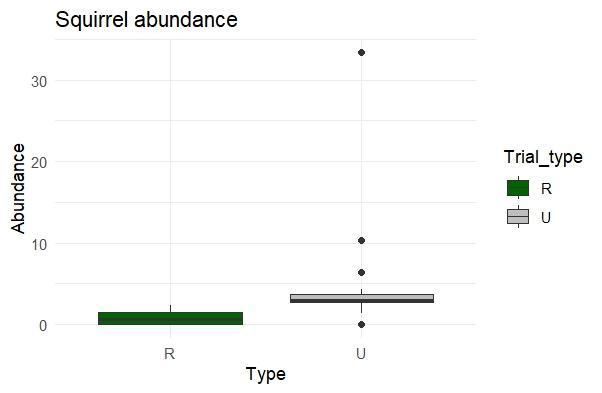

t = -4.1294,

**p-value = 0.00017**

Fig. S5. Abundance of squirrels in the area(within 50m) of the experiment across rural and urban trials

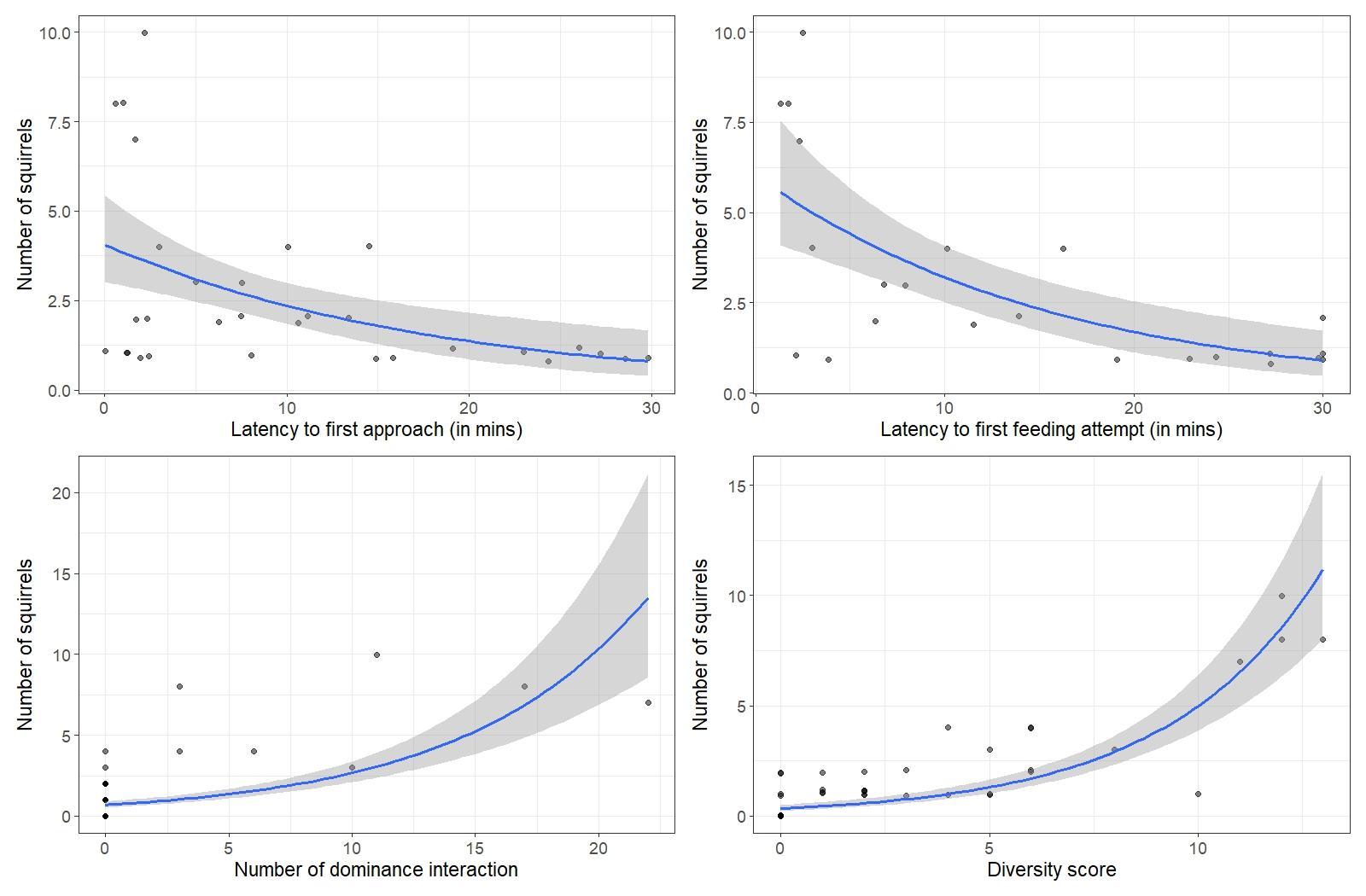

Fig. S6. Effect of squirrel in vicinity(1m) on latency to first approach

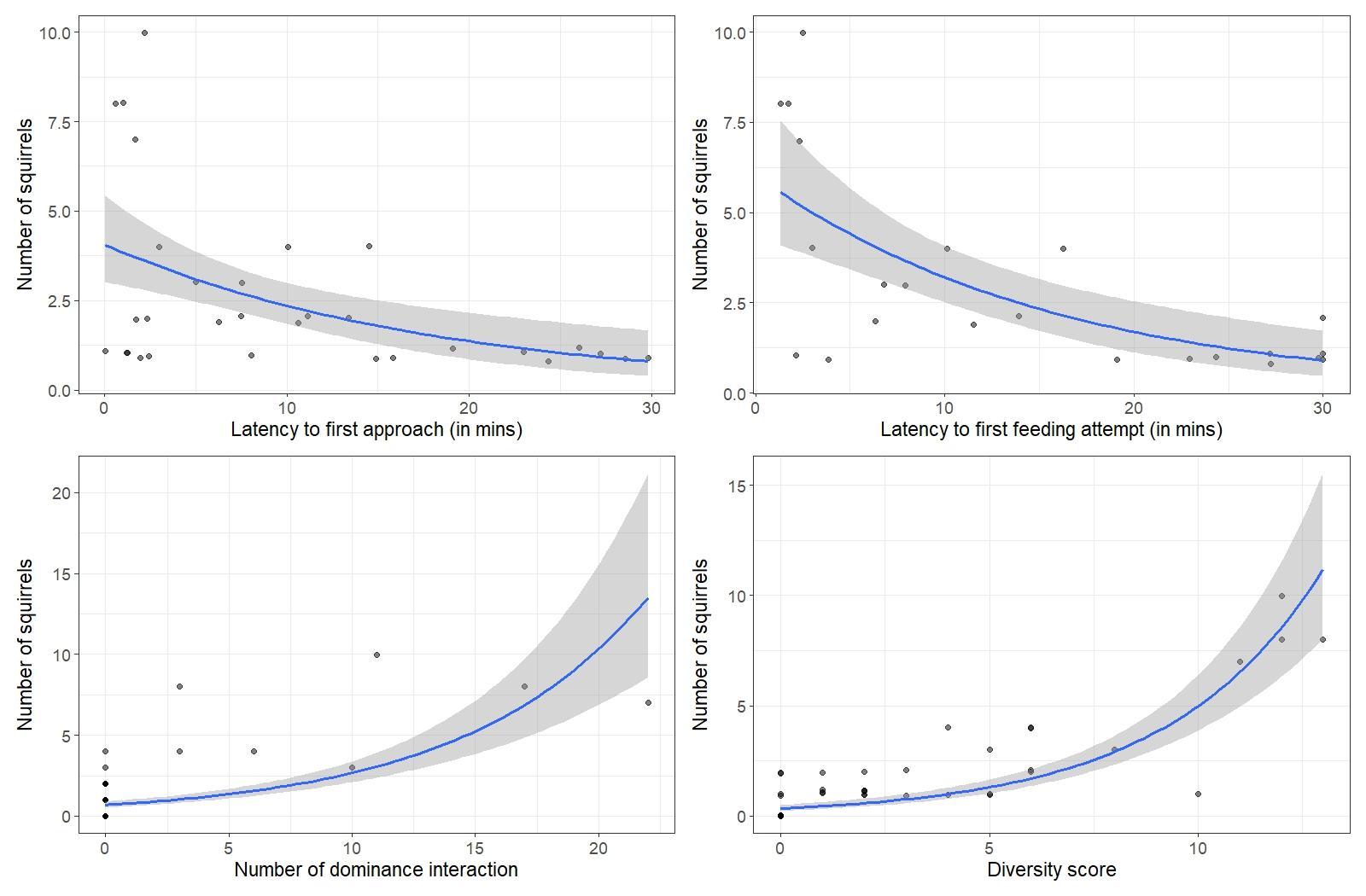

Fig. S7. Effect of squirrel in vicinity(1m) on latency to first feeding attempt

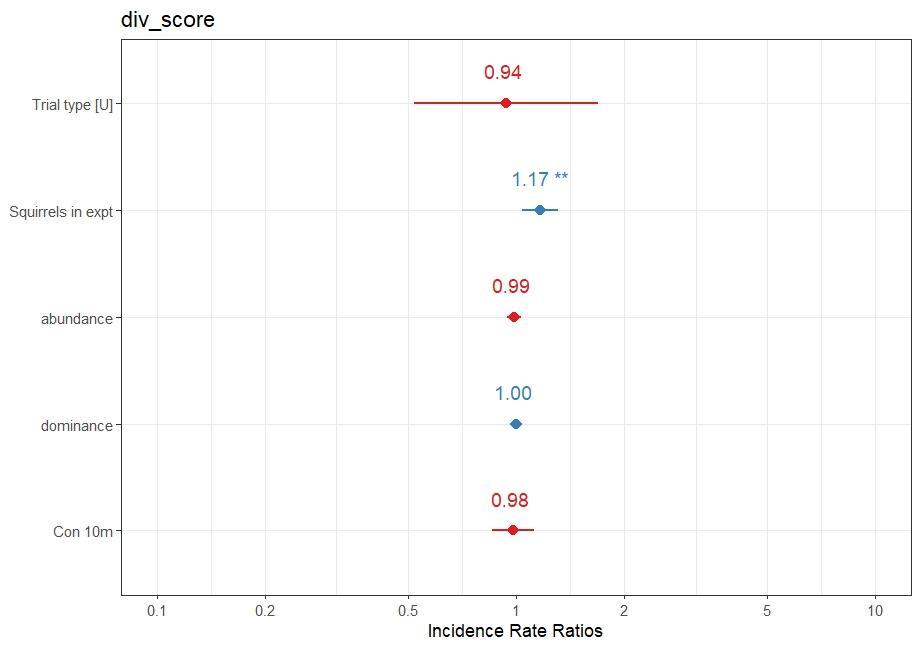

Fig. S8. Effect sizes of factors influencing diversity score

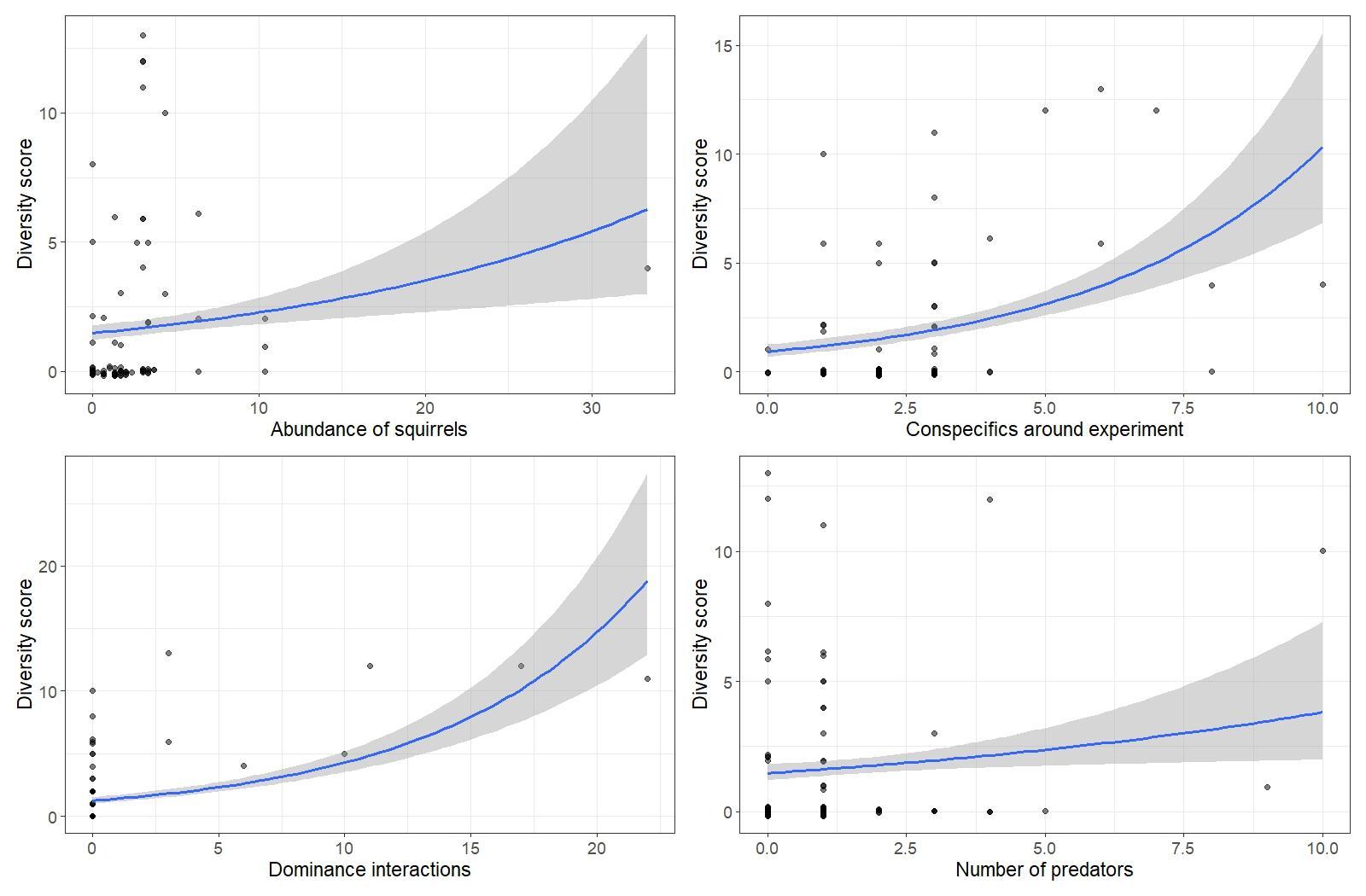

Fig. S9. Regression model of conspecifics in the vicinity(1m) of the experiment on the diversity score

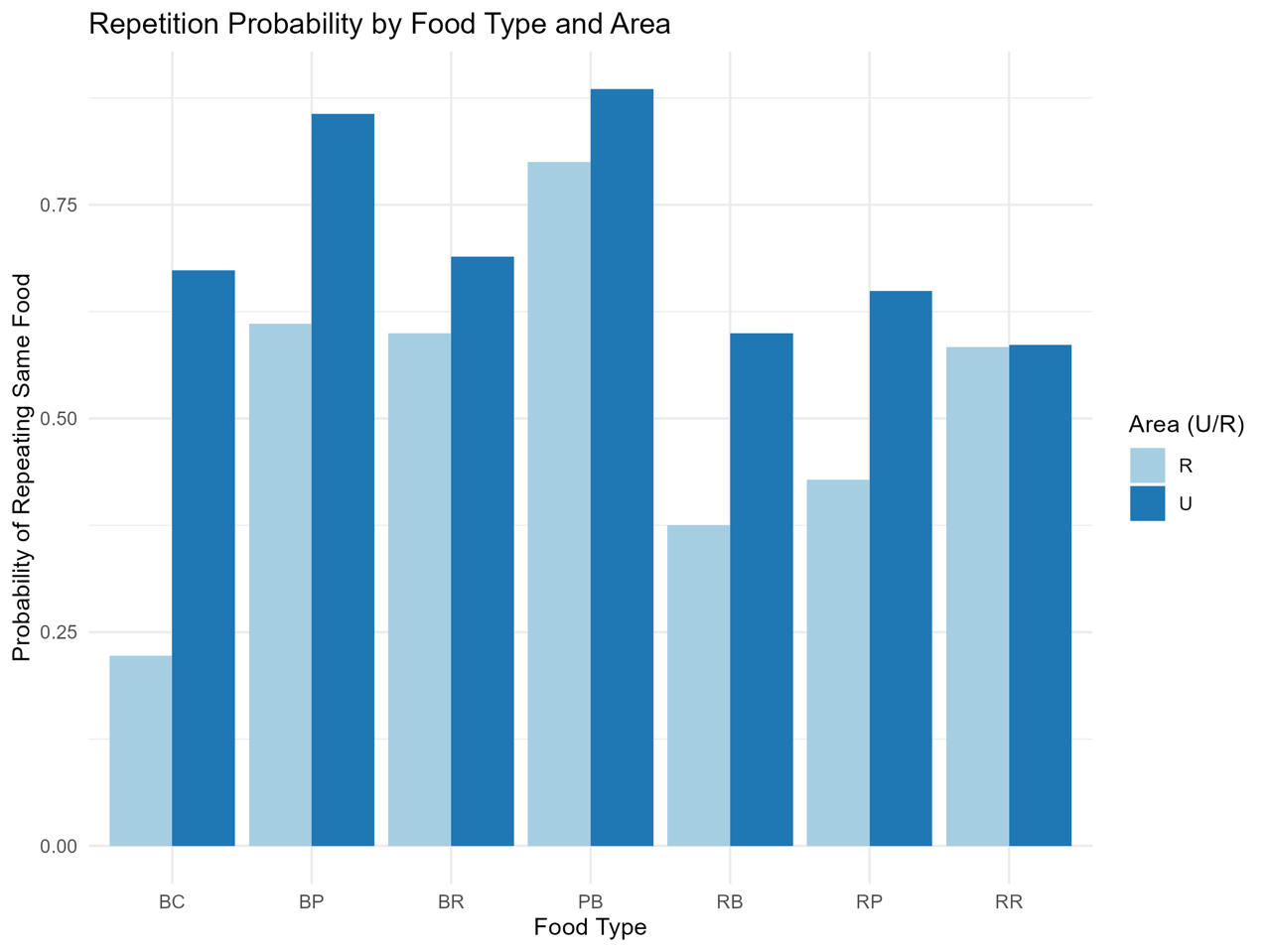

Fig. S9. Bar plot of repetition probabilities of food type in urban and rural areas

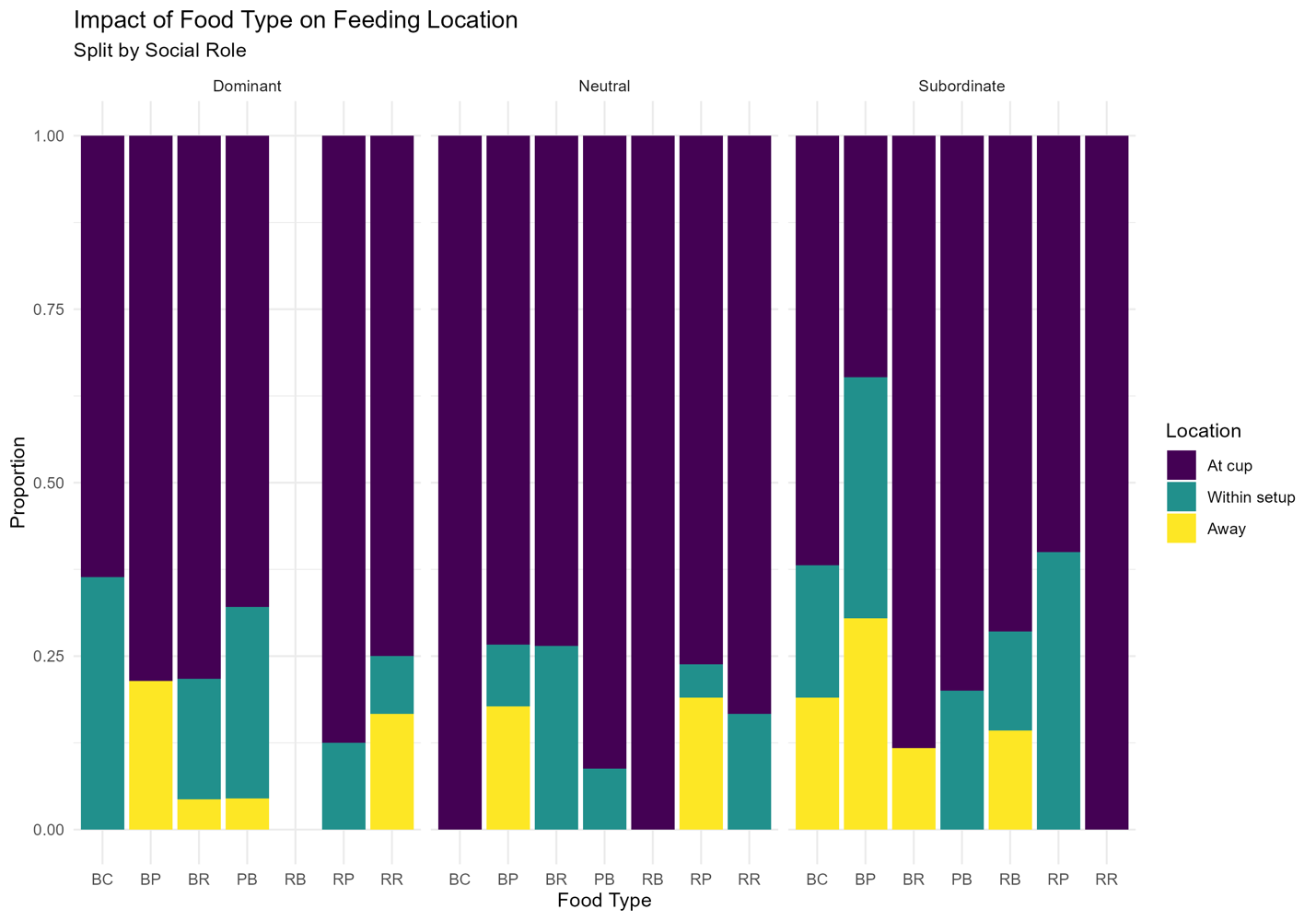

Fig. S11. Bar plot for the effect of food type and feeding location across social roles

Table S1. GLMM model of factors influencing the time taken to first approach

|  | Time taken to first approach | | |
| --- | --- | --- | --- |
| Predictors | Incidence Rate Ratios | CI | P |
| (Intercept) | 35.72 | 12.57 – 101.52 | **<0.001***** |
| Trial type [Urban] | 0.89 | 0.63 – 1.26 | 0.513 |
| Squirrels in vicinity (1m) | 0.64 | 0.54 – 0.75 | **<0.001***** |
| Squirrel abundance | 1.02 | 1.00 – 1.05 | 0.060 |
| Predator presence | 0.69 | 0.50 – 0.94 | **0.018*** |
| Heterospecific presence | 0.88 | 0.57 – 1.35 | 0.558 |
| Conspecifics in 10m | 1.21 | 1.07 – 1.37 | **0.003**** |
| Conspecifics in 50m | 0.92 | 0.83 – 1.02 | 0.105 |
| Random Effects | | | |
| σ^2^ | 0.15 | | |
| τ_00_ | 1.00 _food_appr_first_ | | |
| ICC | 0.87 | | |
| N | 7 _food_appr_first_ | | |
| Observations | 25 | | |
| Marginal R^2^ / Conditional R^2^ | 0.553 / 0.941 | | |

Table S2. GLMM model of factors influencing the time taken to the first feeding attempt

|  | Time taken to first feed | | |
| --- | --- | --- | --- |
| Predictors | Incidence Rate Ratios | CI | p |
| (Intercept) | 28.96 | 10.11 – 82.96 | **<0.001***** |
| Trial type [Urban] | 1.67 | 1.03 – 2.72 | **0.039*** |
| Squirrels in vicinity (1m) | 0.64 | 0.56 – 0.74 | **<0.001***** |
| Squirrel abundance | 1.03 | 1.01 – 1.05 | **0.013*** |
| Predator presence | 0.50 | 0.36 – 0.69 | **<0.001***** |
| Heterospecific presence | 0.79 | 0.51 – 1.21 | 0.278 |
| Conspecifics in 10m | 1.07 | 0.93 – 1.23 | 0.346 |
| Conspecifics in 50m | 1.01 | 0.89 – 1.15 | 0.868 |
| Random Effects | | | |
| σ^2^ | 0.12 | | |
| τ_00_ | 1.00 _food_fed_first_ | | |
| ICC | 0.89 | | |
| N | 7 _food_fed_first_ | | |
| Observations | 22 | | |
| Marginal R^2^ / Conditional R^2^ | 0.544 / 0.950 | | |

Table S3. GLMM model of factors influencing the diversity score

|  | | Diversity score | | |
| --- | --- | --- | --- | --- |
| Predictors | | Incidence Rate Ratios | CI | p |
| (Intercept) | | 3.99 | 2.33 – 6.84 | <0.001 |
| Trial type [U] | | 0.94 | 0.52 – 1.69 | 0.826 |
| Squirrels in vicinity (1m) | | 1.17 | 1.04 – 1.31 | 0.008 |
| Abundance | | 0.99 | 0.94 – 1.03 | 0.506 |
| Dominant interactions | | 1.00 | 0.96 – 1.04 | 0.961 |
| Conspecifics within 10m | | 0.98 | 0.86 – 1.12 | 0.759 |
| Random Effects | | | | |
| σ^2^ | 0.15 | | | |
| τ_00_ _food_fed_first_ | 0.00 | | | |
| N _food_fed_first_ | 4 | | | |
| Observations | 22 | | | |
| Marginal R^2^ / Conditional R^2^ | 0.542 / NA | | | |

Table S4. Ethogram for classification of behaviour for annotation in BORIS software

| **Behaviour** | **Description** | **Modifier** | **Modifier description** | **State/ Event** |
| --- | --- | --- | --- | --- |
| Feeding | Consuming food items: Including nibbling on food items or licking food items held in paws | Feeding directly from the cup | Feeding on the item directly from the experimental cup | S |
|  |  | Feeding from hand | Taking an item from the cup with the hands and then feeding it out of them |  |
|  |  | Position of squirrel while feeding: Sitting | Squirrel is sitting on its hind limbs and feeding from hands/sitting on all fours and feeding directly from a cup |  |
|  |  | Position of squirrel while feeding: Lying | Squirrel holding on to the surface with its hind legs, lying flat with its stomach touching the surface, and feeding from its hands |  |
|  |  | Position of squirrel while feeding: Standing | Squirrel standing on hind limbs and feeding from hands |  |
|  |  | Location of the squirrel while feeding: At the cup | Squirrel either feeds directly from a cup or takes food in its hand and feeds near the cup of the respective food item being consumed |  |
|  |  | Location of the squirrel while feeding: within the setup | Squirrel taking an item from the cup and then feeding within the setup (up to 1-2m) or feeding near a cup other than that of the food item it picked. |  |
|  |  | Location of the squirrel while feeding: Away from the setup | Squirrel taking an item from the cup and moving farther than 2m away from the setup to feed |  |
| Foraging | Looking or searching for food: Includes a series of activities where the face of the individual is towards the surface with multiple bouts of movement, and occasional sniffing/handling. | NA | NA | S |
| Moving | moving from one place/ position to another at a slow/moderate speed on a surface | 1. Moving up  2. Moving down  3. Away  4. Towards  5. With food | 1. Slowly moving upwards on a vertical surface  2. Slowly moving downwards on a vertical surface  3. Putting some distance between the focal and others/objects  4. Closing distance between the focal and others/objects  5. Carrying food while moving | S |
| Running | Moving quickly on a surface over a longer distance | 1. Running up  2. Running down  3. With food | 1. Moving quickly upwards on a vertical surface over a longer distance  2. Moving quickly downwards on a vertical surface over a longer distance  3. Carrying food while running | S |
| Approach experiment | Entering the vicinity(1-2m from the setup) and actively moving towards/approaching the experimental setup |  | Stop and interact with the whole cafeteria experiment | S |
| Approach item | Stop and interact with a particular food choice from within the experiment | Food choices | 7 food choices - PB, RR, BR, RB, BC, RP, BP | E |
| Sniffing | Approaching an object, taking the nose close to it, and twitching the nose | Food choices | 7 food choices - PB, RR, BR, RB, BC, RP, BP | S |
|  |  | Sniffing item | Approaching a particular food choice within the experiment, taking the nose close to it, and twitching the nose |  |
| Chasing/Displacing | Squirrel(s) running behind other squirrel(s)/ a squirrel taking the place of another | Individual  1. Chaser  2. Chasee  3. Supplanter  4. Supplantee | 1. Focal squirrel is chasing other individual(s)  2. Focal squirrel is being chased by other individual(s)  3. Focal squirrel (momentary movement) displaces other individuals  4. Focal squirrel is (momentary movement) displaced by another individual | S |
|  |  | Direction  1. Stay  2. Away  3. Towards | 1. Focal squirrel stays within the setup  2. Focal squirrel moves away from the setup  3. Focal squirrel moves towards the setup | S |
|  |  | Food  1. With food  2. Without food | 1. Carrying food while chasing/being chased  2. Not carrying food while chasing/being chased | S |
| Lying  (excludes feeding) | Flat posture, stomach resting, touching the surface | 1. Resting  2. Frozen | 1. Lying where limbs and tail are relaxed.  2. Lying without any movement on a surface for a time period. Usually occurs when a threat is present or alarm calls are heard. | S |
| Sitting  (excludes feeding) | Posture/ position while not moving on a horizontal surface | 1. On hind limbs  2. On fours | 1. Vertical posture, body resting on hind limbs that are on a horizontal surface  2. Sitting on all fours on a horizontal surface (stomach not touching surface) | S |
| Standing  (excludes feeding) | Stretching upwards with hind limbs on a horizontal surface |  |  | S |
| Calling | Producing sound in patterns | 1. Alarm call  2. Mating call  3. Contact call | 1. Sound produced in a specific pattern to alert other squirrels  2. Sound produced in a specific pattern to attract mates | S |
| Alert | Looking for signs of danger in the surrounding environment |  |  | S |
| Searching | Moving slowly on the ground, head turning from one side to another at least at a 1m distance away from any food source (natural/experimental setup) |  |  | S |
| Leaping | Launching forward from one location by pushing on hind legs, covering a distance and landing, forelimbs first, at another location. This behaviour occurs independently of a chase sequence. | 1. With food | Focal squirrel carries food with it as it leaps | E |
| Mating chase | Squirrels initiating and partaking in chases during a mating event | 1. Female-male chase  2. Male-male chase | 1. Female is chased by a male to initiate copulation.  2. Multiple males initiate a chase to interrupt and drive away a copulating male | S |
| Mating | Two individuals are in a copulatory position (usually stationary) - male mounting a female |  |  | S |
| Collecting nest material | Squirrel looking for, handling, and carrying non-food items |  |  | S |
| Grooming | Cleaning/sniffing | 1. Autogrooming  2. Allogrooming | 1. Grooming self - includes sniffing, cleaning  2. Grooming conspecifics | S |
| Upturn  (excluding during chases) | Quickly overturn an object while moving past it/handling it |  |  | E |
